# Mechanical licensing of functional dendritic cell states for enhanced T cell priming

**DOI:** 10.64898/2026.05.19.725170

**Authors:** Yu-Chang Chen, Amanda S. Bluem, Fatemeh Tavakoli Joorabi, Kexin Zhang, Nghi M. Tran, Shuchen Zhang, Hardik Makkar, Kyle H. Vining

## Abstract

The plasticity of dendritic cell (DC) functional state is a major hurdle in DC therapy, yet how DCs acquire distinct states independent of ontogeny remains poorly understood. Here, we demonstrate that changes in matrix stress relaxation mechanically educate DCs to adopt distinct, persistent functional states even after the removal of mechanical cues. Stem cell-derived DCs cultured in a fast-relaxing environment exhibited enhanced antigen presentation, faster migration, and higher expression of T cell-recruiting chemokines. Slow-relaxing DCs, biased towards pro-inflammatory cytokine secretion, were enriched for gene signatures associated with lipid accumulation and stress response. These mechanical responses were conserved across human and murine DCs. Using ovalbumin (OVA) as the model antigen, fast-relaxing DCs elicited a CD8+-biased response in vitro, with higher antigen-specific CD8+ T cell activation and proliferation. In vivo adoptive cell transfer of mechanically educated DCs demonstrated that the fast-relaxing matrix licensed DCs to induce a potent draining lymph node T cell response with more antigen-specific T cells and higher restimulation potential. We further showed that DCs sensed matrix stress relaxation through PI3K signaling and actin branching, mediated by the concerted signaling of IL-4 and GM-CSF. Together, these findings demonstrate the role of matrix stress relaxation on the functional state of DCs and suggest a novel approach to enhance ex vivo cellular engineering by targeting mechanical signaling.

**Graphical Abstract:** Stem cell-derived dendritic cells (DCs) generated ex vivo are engineered using biomaterial platform with tunable matrix stress relaxation. Mechanical education of DCs is licensed by cytokine signaling, actin branching, and PI3K signaling. Fast-relaxing DCs exhibit higher antigen presentation and faster migration, which enhances their capacity to prime and activate antigen-specific CD8+ T cells.

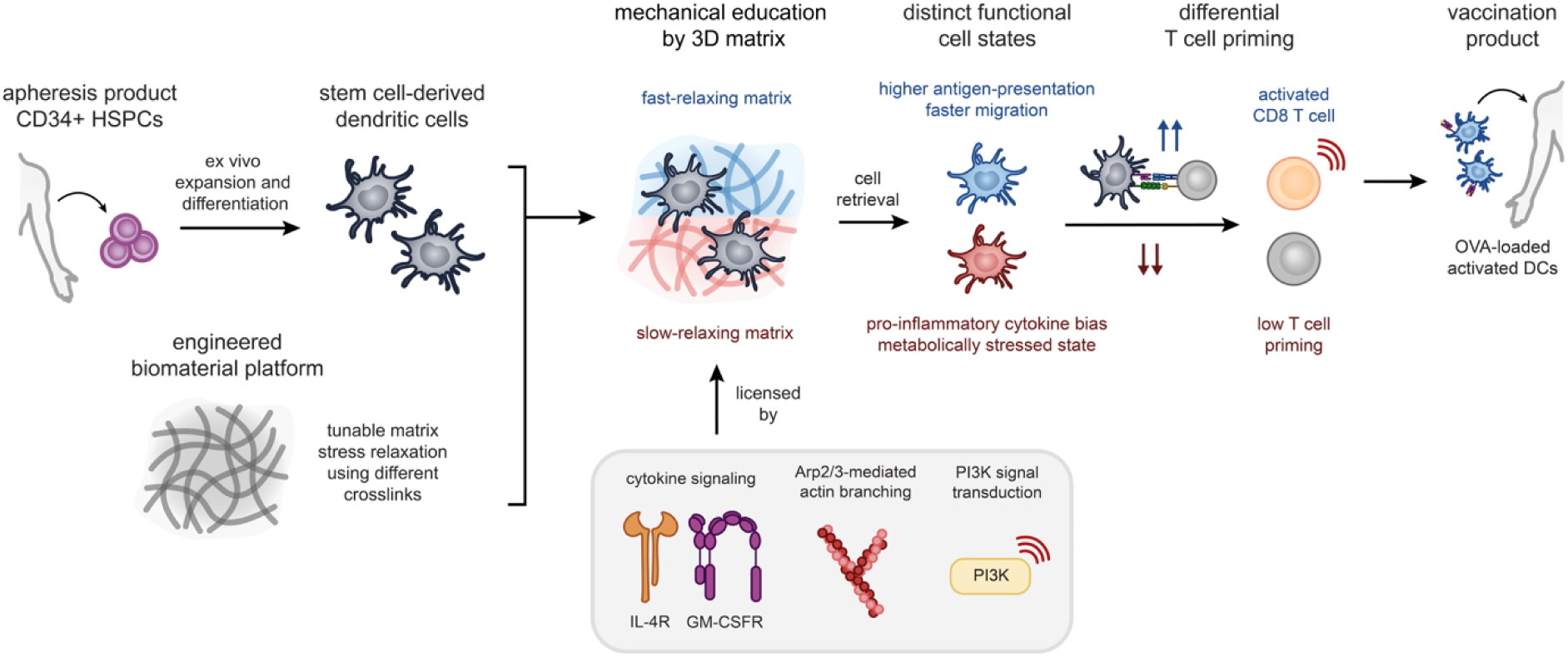

## Introduction

Harnessing the functions of dendritic cells (DCs) for immunotherapies has become a rapidly expanding strategy to treat cancer and autoimmune diseases, given their ability to shape the adaptive immune response. However, a lack of response or an opposite response from DC therapies in clinical trials is hindering their therapeutic efficacy^1–4^. One underlying mechanism is the profound complexity of myeloid heterogeneity and its plasticity^5–8^. Upon interacting with factors in the tissue microenvironment, they take on different functional states and drive differential immune responses regardless of their ontogeny, often contributing to disease progression^9^. For example, both cDC1 (conventional DC) and cDC2 adopt a mature regulatory state after tumor antigen uptake to influence anti-tumor immunity^10^. Although research in this area has received significant attention, little is known about how DCs acquire distinct functional states in distinct microenvironments. Existing work identified biochemical cues, such as immune checkpoints, cytokine signaling, oxidative species, and tumor-derived factors, are capable of shaping cell state changes^5–9^. We hypothesize that physical cues, such as stress relaxation of the extracellular matrix (ECM), also play a role in reprogramming the functional state of DCs and can be exploited for cellular engineering.

The field of mechanobiology has established that cells respond to their mechanical environment and permanently alter their cell state through epigenetic modificiations^11–13^. Studies done on immune cells, with some specifically on DCs, show that they are sensitive to changes in the stiffness and viscoelasticity of their surrounding ECM^14–17^. Cellular response to stiffness has been under active investigation, whereas viscoelastic properties, such as matrix stress relaxation, have been understudied. Since matrix stress relaxation is a dynamic property with physiological relevance in remodeled tissues, we are interested in investigating its effect on DC fate using a biomaterial platform with tunable stress relaxation. Nevertheless, analyses in traditional mechanobiology studies are often performed while cells remain exposed to mechanical cues.

To demonstrate that cell-state changes due to mechanical regulation persist after mechanical cues are removed, we performed all of analyses on DCs retrieved from hydrogels and reseeded onto tissue culture plastic. We term this “mechanical education,” where DCs remember being in distinct mechanical environments, causing them to have a permanent change in cell state and carry out different immune functions. In brief, we showed that DCs educated by the fast-relaxing matrix were licensed for enhanced T cell priming both in vitro and in vivo. Our results suggest a novel approach for ex vivo cellular manufacturing that target the mechanical axis to enhance the efficacy of cell products, bridging basic science to clinical translation^18^. Moreover, the versatility of biomaterial platforms allows for additional biochemical molecules to be presented to the cells, thereby synergistically activating them through both biochemical and mechanical cues.

## Results

### Human stem cell-derived HLA-DR+ CD11c+ dendritic cells (DCs) are mechanically educated in viscoelastic extracellular matrix (ECM) hydrogels

We have developed a biomaterial strategy for generating DCs from autologous stem cell sources for therapeutic applications. Human DCs are traditionally differentiated from peripheral blood-derived monocytes or isolated by magnetic selection from blood mononuclear cells. However, DCs are less than 1% of mononuclear cells in the peripheral blood^19,20^, and monocyte-derived DCs (moDCs) are limited by their distinct ontogeny compared to naïve stem cell-derived DCs^21,22^. Recent advances have shown that DCs differentiated from CD34+ hematopoietic stem and progenitor cells (HSPCs) in vitro resemble the functions and transcriptome of blood DCs from the same subset^23,24^. In addition, CD34+ HSPCs expand to large numbers while maintaining their self-renewal and differentiation capacity in vitro using cytokine and small molecule cocktails^25–27^. First, we showed that large numbers of stem cell-derived DCs can be obtained from sequential CD34+ HSPC expansion and differentiation in vitro **(Fig. 1A)**. Peripheral blood G-CSF-mobilized HSPCs showed a larger expansion capacity than inducible pluripotent stem cell (iPSC)-derived CD34+ HSPCs **(Fig. S1A)**. After two expansion passages with IL-3, SCF, TPO, FLT3-L, and UM729, mobilized CD34+ HSPCs expanded to 80-fold while retaining more than 80% of CD34+ population **(Fig. S1B-C)**. These expanded HSPCs were differentiated into HLA-DR+ CD11c+ DCs using FLT3-L and GM-CSF^28–30^ and turned on expression of lineage commitment marker CD1c **(Fig. S1D)**.

**Figure 1.**
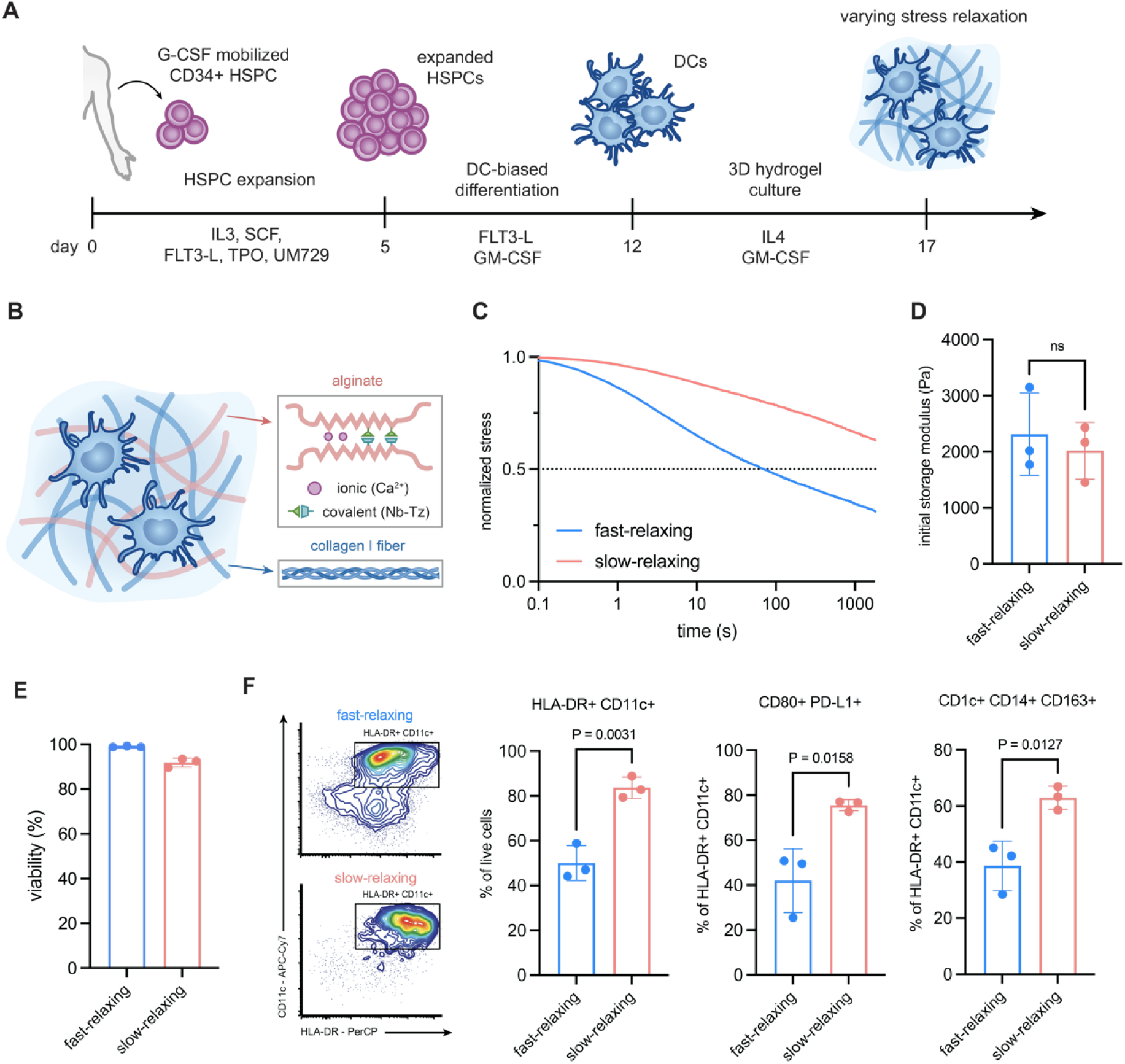
Interpenetrating click-modified alginate and collagen networks tunes matrix stress relaxation to alter the phenotype of stem cell-derived dendritic cells (DCs). **(A)** Workflow to generate ex vivo stem cell-derived DCs. Human G-CSF mobilized CD34+ hematopoietic stem and progenitor cells (HSPCs) were expanded to large numbers and differentiated into DCs using cytokine cocktails. DCs were then encapsulated in 3D hydrogels with tunable stress relaxation. (B) Schematic of the interpenetrating network of alginate (red) and type I bovine telo collagen (blue). Alginate was chemically modified with norbornene (Nb) and tetrazine (Tz), which spontaneously formed covalent crosslinks upon mixing. Unmodified alginate chains formed ionic crosslinks through Ca^2+^ binding. The ratio of modified and unmodified alginate tuned matrix stress relaxation due to the physical differences in their bonding. (C) Stress relaxation test at 10% shear strain showed faster relaxation from the unmodified alginate (blue) compared to the click-modified alginate (red). (D) Rheological characterization of the fast-relaxing and slow-relaxing hydrogels had no statistical difference of storage modulus. (E) DCs retrieved from the 3D hydrogels after 5 days of culture had no impact on cell viability. (F) Representative flow cytometry plots showing the expression of HLA-DR and CD11c on the mechanically primed DCs. Slow-relaxing hydrogels promoted higher surface expression of phenotypic maturation markers, such as HLA-DR, CD11c, CD80, and PD-L1, and were enriched in lineage differentiated CD1c+ CD14+ CD163+ DCs. Data are shown as mean ± SD, and P-values from two-tailed unpaired Student’s t-tests are indicated. Data in C and D are representative of n = 3 independent samples where only one replicate of each is plotted in D. Data in E and F are representative of n = 3 independent biological replicates from different hydrogel samples using the same human donor.

Next, we investigated the role of 3D matrix stress relaxation on the functional state of DCs in interpenetrating network (IPN) hydrogels composed of type I collagen and chemically modified low viscosity alginate^31–33^ **(Fig. 1B)**. Type I collagen (4 mg/mL) was added to the system to provide adhesive cues and mimic physiological 3D tissue environment. Native alginate chains form ionic crosslinks through divalent cation binding at the G-blocks, and the click-modified alginate chains form covalent crosslinks through the Diels-Alder reaction of norbornene (Nb) and tetrazine (Tz) conjugated on the alginate backbone **(Fig. S2A)**. The Nb-Tz alginate network showed slower stress relaxation and lower tan(delta) viscoelastic properties due to the physical differences between the ionic and covalent crosslinks **(Fig. 1C, S2C)**. Therefore, the rate of stress relaxation can be tuned by the relative mode of crosslinks of the gel formulation independent of adhesive ligand density and pore size. Previous studies have shown that DCs are sensitive to the stiffness of their surroundings^16,34^, so we optimized the formulation to achieve a matched initial storage modulus **(Fig. 1D)**. Stem cell-derived DCs were encapsulated in these hydrogels supplemented with IL-4 and GM-CSF to promote further differentiation and maintain viability^35^ **(Fig. 1E)**. After 5 days of 3D culture, dendritic cells educated by the slow-relaxing matrix showed an increased frequency of HLA-DR+ CD11c+ population expressing maturation markers CD80, PD-L1, and CCR7 **(Fig. 1F, S1E)**. Moreover, the slow-relaxing DCs were polarized into the CD1c+ CD14+ CD163+ subset that is associated with inflammation^36,37^. These observations indicate that matrix stress relaxation serves as a mechanical cue to govern the phenotypic states of DCs.

### Mechanically educated DCs exhibit functional differences after TLR activation

Next, we examined whether matrix stress relaxation affected the functional state of dendritic cells and whether these states persisted after retrieval from 3D ECM gels. Myeloid cells adopt distinct cell states in response to environmental factors due to their plasticity, and different cell states carry out different immune functions^8,9,38^. We retrieved DCs from the hydrogels and reseeded them in tissue culture plastic or collagen gel for functional analyses **(Fig. 2A)**. From the fast-relaxing matrix, immature DCs exhibited a higher 3D migration rate in collagen gels with no changes in phagocytic capacity compared to those retrieved from the slow-relaxing matrix **(Fig. 2B, S3A)**. Upon TLR3 activation, mechanically educated DCs expressed a distinct ligand profile where slow-relaxing DCs upregulated CD86 and CCR7, and fast-relaxing DCs upregulated CD80 and checkpoint ligand PD-L1 **(Fig. 2C, S3B)**. Slow-relaxing DCs exhibited a pro-inflammatory bias with higher secretion of IL-6, TNF-α, and IL-1β, whereas the fast-relaxing DCs secreted more anti-inflammatory IL-10 **(Fig. 2D, S3C)**. The pro-inflammatory bias of slow-relaxing DCs was independent of the type of TLR agonist **(Fig. S4)**. Mixed lymphocyte reaction involves the co-culture of allogeneic T cells and antigen-presenting cells, which provides an approximate measure of DCs’ ability to activate T cells through MHC-mismatch^39,40^. Surprisingly, the slow-relaxing DCs, which express high levels of co-stimulatory ligands and pro-inflammatory cytokines, produced a weaker allogeneic T cell response **(Fig. 2E)**. These findings suggest that slow-relaxing educated DCs resemble hyper-inflammatory DCs, similar to the myeloid cells observed in chronic inflammation characterized by a pro-inflammatory cytokine signature and low antigen presentation^41^. Conversely, the ability to prime T cells is enhanced in fast-relaxing DCs. These data show that matrix-educated DCs adopt distinct functional states even after mechanical cues are removed, indicating a mechanical role of myeloid plasticity. On the contrary, proper regulation of immune activation in fast-relaxing DCs enhances their ability to prime T cells. In both cases, changes in stress relaxation educated DCs to adopt distinct functional states even after mechanical cues were removed, indicating a novel mechanism of myeloid plasticity.

**Figure 2.**
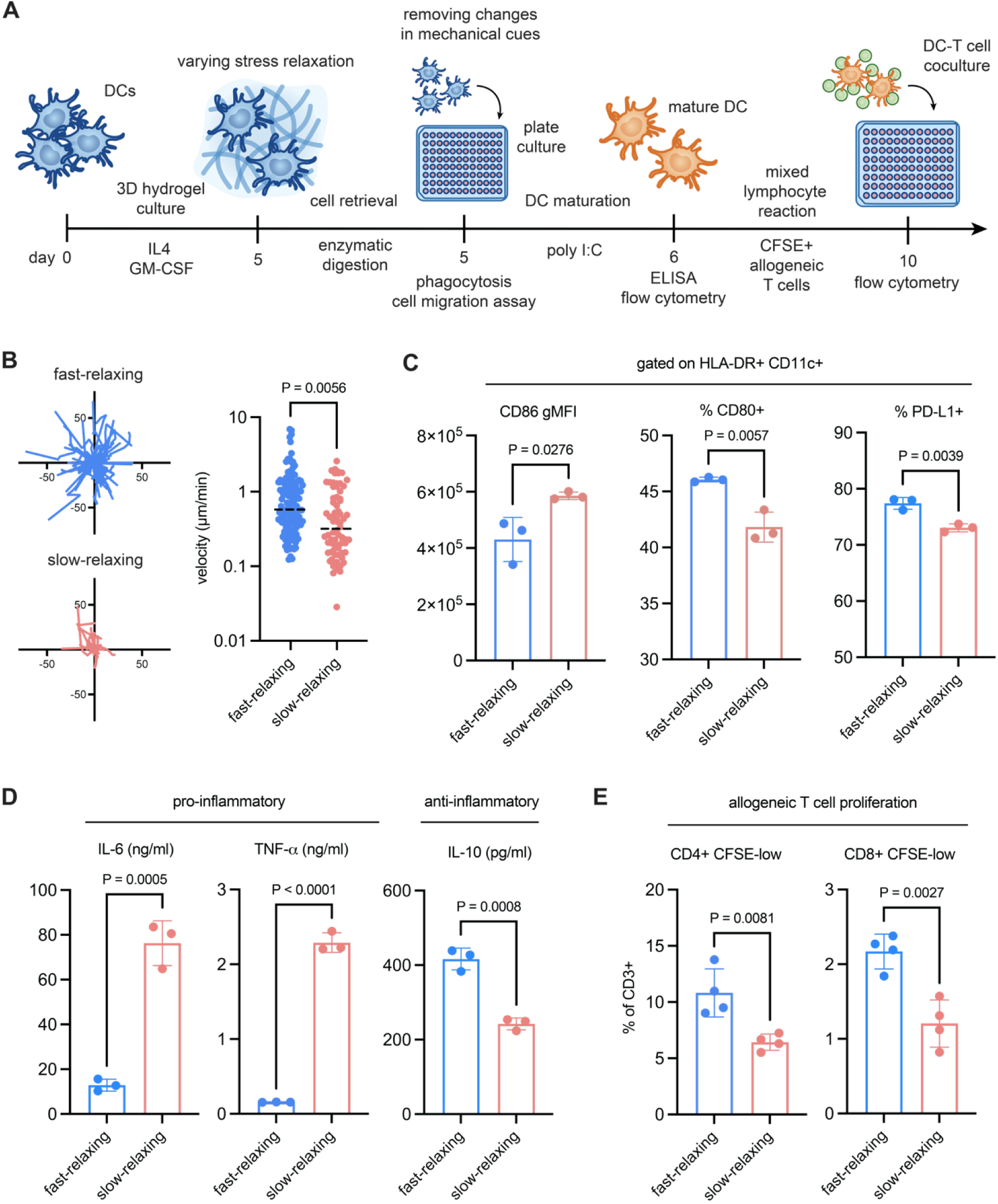
Matrix stress relaxation educates stem cell-derived DCs and alters their immune functions. **(A)** Workflow to assess the functional outputs of mechanically educated DCs. DCs were cultured in 3D hydrogels with varying stress relaxation for 5 days and retrieved through enzymatic digestion of the matrix. Retrieved cells were reseeded in tissue culture plastic for downstream functional analyses. **(B)** Spider plots of migrating DCs in 1mg/ml type I collagen hydrogel over 40 minutes. DCs retrieved from the fast-relaxing hydrogel had a higher migration rate despite being reseeded in the same matrix condition. **(C)** Flow cytometry characterization of co-stimulatory and regulatory ligands at the immunological synapse after TLR3 activation of mechanically educated DCs in plate culture. Fast-relaxing hydrogel promoted CD80+ and PD-L1+ DCs but with lower CD86 expression. **(D)** ELISA quantification of secreted cytokines by the mechanically educated DCs. Slow-relaxing matrix promoted a pro-inflammatory bias of heightened IL-6 and TNF-α secretion while reducing level of IL-10. **(E)** Proliferation of allogeneic CD3+ T cells stained with CFSE after co-culture with mechanically educated DCs. DCs retrieved from the fast-relaxing matrix promoted higher frequency of CFSE-low cells, indicating higher proliferation of both CD4+ and CD8+ allogeneic T cells. Each dot in B represents a single cell, and P-value from two-tailed unpaired Student’s t-tests is indicated. Data in C, D, and E are shown as mean ± SD, and P-values from two-tailed unpaired Student’s t-tests are indicated. Data are representative of biological replicates from different hydrogels using the same human donor, and each dot in the bar charts indicates one replicate.

### Fast-relaxing hydrogels support a T cell-priming transcriptomic state in DCs

Next, we performed bulk RNA sequencing of DCs isolated from hydrogels to evaluate their transcriptome prior to TLR activation. Principal component analysis (PCA) revealed that the slow-relaxing DCs harbored a distinct transcriptomic state compared to the fast-relaxing and the plate culture DCs **(Fig. 3A)**. Fast-relaxing DCs upregulated genes that are essential for T cell priming, such as HLA family (HLA-A, HLA-DPB1, HLA-DRB1) and T cell chemokines (CCL5, CCL18, CCL23) **(Fig. S5A)**. Pathway analysis confirmed the observation with upregulation of pathways including MHC assembly, T cell cytokine production, and T cell activation **(Fig. 3B)**. Conversely, pathways related to mechanical stimulus, inflammatory response, and PI3K signal transduction were downregulated. Heatmap analysis of all genes in the HLA family showed that fast-relaxing DCs had a higher expression of antigen presentation machinery **(Fig. 3C)**. Consistent with the ELISA result, mechanically educated DCs had a bias towards their cytokine and chemokine response **(Fig 3D)**. Slow-relaxing DCs exhibited a pro-inflammatory bias with increased expression of IL6, IL1B, and TNF, and a pro-tumor state with higher TGFB1 and CCL22^42,43^. On the other hand, fast-relaxing DCs were enriched in anti-inflammatory cytokine IL10 and T cell-recruiting chemokines CCL5, CCL17, CCL18, and CXCL10. Similar to the different expression of cytokines, co-stimulatory molecules were differentially enriched in the mechanically educated DCs **(Fig. S5B)**. Notably, transcription factors such as BATF3, RELB, and NFKB1 that drive DC maturation were upregulated in fast-relaxing DCs^44–46^. The increased expression of regulatory programming genes SOCS1 and SOCS3 in fast-relaxing DCs supported our hypothesis that the slow-relaxing DCs were overactivated due to a loss of checkpoint mechanisms^47,48^ **(Fig. S5C)**. Lipid accumulation and dysregulated glycolysis lead to dendritic cell dysfunction and impair their ability to activate T cells^49,50^. Slow-relaxing DCs upregulated expression of hallmark genes related to lipid metabolism and accumulation (ABCG1, CD36, PPARG, MSR1, APOE), which has been shown to associate with a stress response in DCs^51^, indicating a potential reason for their hyper-inflammatory state **(Fig. 3E)**.

**Figure 3.**
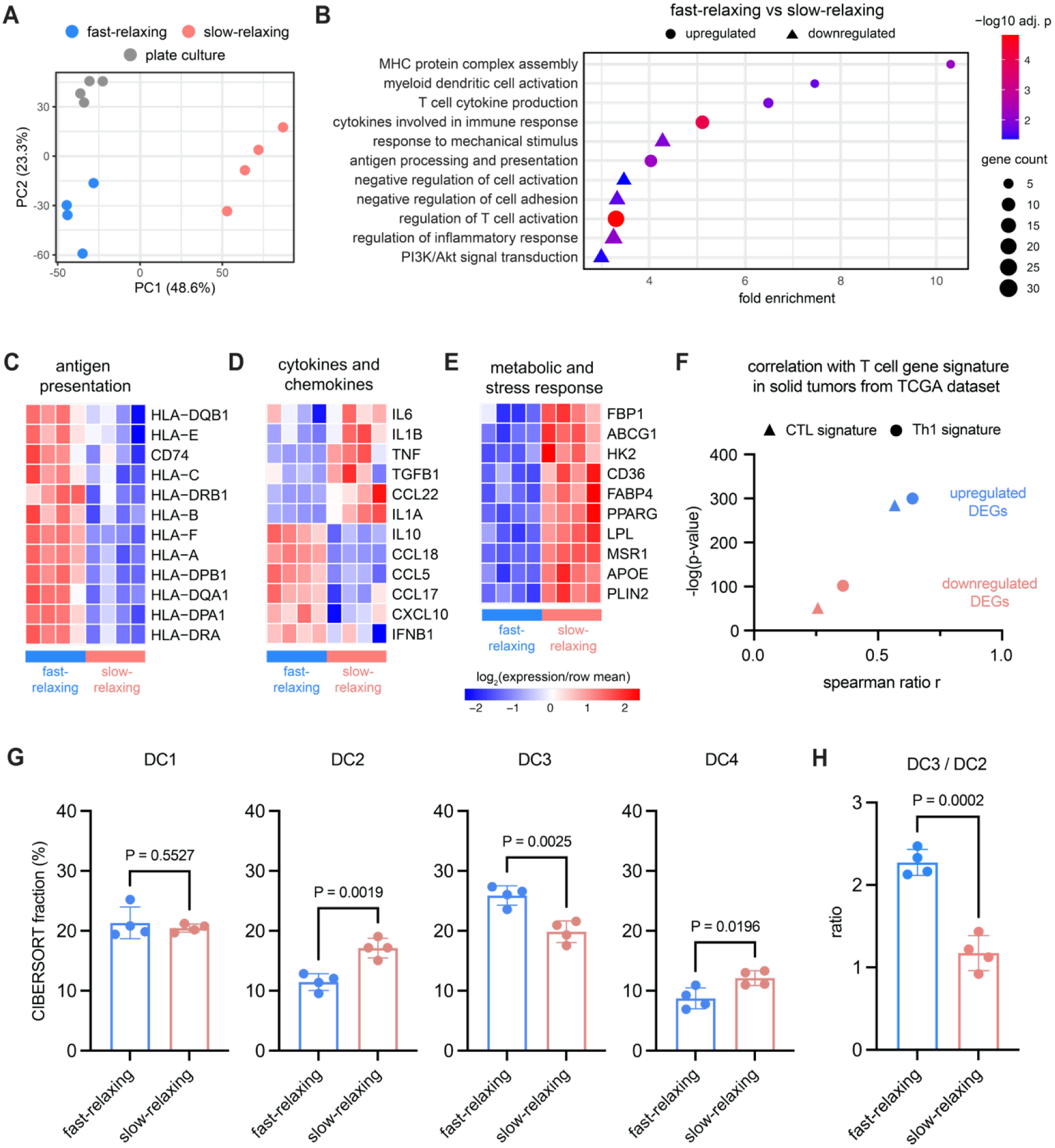
Transcriptomic analysis reveals distinct DC states primed by the relaxation rate of the matrix. **(A)** Principal component analysis (PCA) generated three clusters of DCs. The major PC (48.6%) separated the slow-relaxing condition from the fast-relaxing condition and the plate culture control. **(B)** Gene Ontology (GO) enrichment of differentially regulated pathways in fast-relaxing versus slow-relaxing environment. The bubble plot showed upregulation of pathways related to antigen presentation and T cell activation, and downregulation of pathways related to inflammation and PI3K/Akt signal transduction. Bubble size indicates the gene count; bubble shape indicates down- or up-regulation; bubble color indicates the adjusted P-value of the GO analysis. **(C, D, E)** Heatmaps of genes associated with antigen presentation, cytokines, and metabolic stress response. The fast-relaxing matrix promoted antigen presentation and biased towards a T cell-recruiting cytokine profile, whereas the slow-relaxing matrix induced a lipid accumulation state and a pro-inflammatory cytokine profile. **(F)** Spearman correlation of differentially expressed genes (DEGs) with cytotoxic T lymphocyte (CTL) and T helper 1 cells (Th1) gene signature in The Cancer Genome Atlas (TCGA) cohort. The cohort consists of major solid tumor types including BRCA, COAD, HNSC, LUSC, and SKCM (n = 3044). Enrichment score per sample was calculated using ssGSEA, and spearman ratio r was plotted against the - log(p-value) of the correlation. The upregulated DEGs showed higher correlation with both T cell signatures in solid tumors. **(G)** CIBERSORT deconvolution revealed that matrix stress relaxation regulated DC subset bias. The slow-relaxing matrix induced a higher frequency of the monocyte-like DC4 subset, **(H)** whereas the fast-relaxing matrix polarized the conventional CD1c+ DC subset into the more immunogenic state (DC3). Data in A are representative of n = 4 biological replicates from different hydrogels using the same human donor. Each column of the heatmap in C, D, and E indicates a single biological replicate. Each dot in G and H indicates one replicate, and are shown as mean ± SD. P-values from two-tailed unpaired Student’s t-tests are indicated.

We asked whether these observed transcriptomic changes between mechanically educated DCs relate to T cell recruitment in solid tumors. Gene expression data of five solid tumor types were retrieved from TCGA, and enrichment scores of the differentially expressed genes (DEGs) and two T cell gene signatures were calculated for each sample. Spearman correlation between DEGs and the T cell gene signatures was performed, and the Spearman correlation coefficient, which measures the strength of the correlation, was obtained **(Fig. S5D)**. Both the cytotoxic T lymphocyte (CTL) and T helper 1 cell (Th1) signatures had higher spearman ratio values with the upregulated DEGs, indicating that the fast-relaxing DCs had a stronger correlation with effector T cell infiltration in the tumors **(Fig. 3F)**. Given that these transcriptomic changes suggested distinct cell states in mechanically educated DCs, we sought to determine whether these states correspond to changes in DC subsets. DCs, like other myeloid cells, are heterogeneous in nature, where different subsets carry varying functions. Single-cell RNA sequencing of peripheral blood DCs identified six DC subsets based on their transcriptome^39^. We applied CIBERSORT deconvolution to bulk RNA sequencing data from mechanically educated DCs, using the peripheral blood DC data set (GSE94820) as a reference to measure the relative abundance of subsets based on transcriptomic data. The quality of the analysis was validated by verifying that there were no statistical differences between the correlation of each sample with the data set and the absolute total scores **(Fig. S5E)**. We found that the fast-relaxing DCs had a higher frequency of the inflammatory CD1c+ DC3 subset, whereas the slow-relaxing DCs were biased towards the conventional CD1c+ DC2 subset **(Fig. 3G, H)**. The two subsets were shown to have similar ability to activate naïve T cells but respond to TLR differently^39^. Moreover, the slow-relaxing DCs were enriched in the monocyte-like DC4 subset that is associated with low capacity to stimulate T cells^39,52^. These results demonstrate that matrix stress relaxation induced a transcriptomic shift in DCs, shaping their cell state and endowing them with a distinct capacity to prime T cells.

### Mechanical education of DCs in stress-relaxing hydrogels is consistent across human and murine DCs

We generated murine stem cell-derived DCs to assess whether the effect of matrix stress relaxation is species-dependent. Murine cKit+ HSPCs were isolated from long bones (C57BL6 mice) and differentiated with mouse FLT-3L and GM-CSF^53–55^ **(Fig. 4A)**. Murine DCs (mDCs) were then encapsulated in the fast- and slow-relaxing hydrogel formulations and retrieved for analysis. Consistent with findings in human DCs, slow-relaxing mDCs had a higher frequency of the I-A/I-E+ CD11c+ population expressing CD80 and PD-L1 **(Fig. 1F, S6A)**. After TLR activation, the fast-relaxing mDCs also upregulated CD80 and PD-L1 expression compared to slow-relaxing mDCs **(Fig. 2C, 4B, S6B)**. Since mDC activation involved adding OVA protein as the antigen source, mechanically educated mDCs were stained with antibodies that bind the MHC-peptide complex (H2K^b^-SIINFEKL) to assess antigen presentation. Fast-relaxing mDCs had a stronger expression of MHC-peptide complex, suggesting their superior ability to present antigens to T cells **(Fig. 4B)**. In terms of subset distribution, fast-relaxing mDCs were enriched in the SIRPα+ CD8α- cDC2 subset, but no differences were observed in the CD8α+ cDC1 subset **(Fig. S6C)**. Interestingly, the slow-relaxing mDCs mainly consisted of populations lacking markers of both conventional subsets, which agrees with the CIBERSORT deconvolution showing the human slow-relaxing DCs were enriched in the non-conventional DC4 subset.

**Figure 4.**
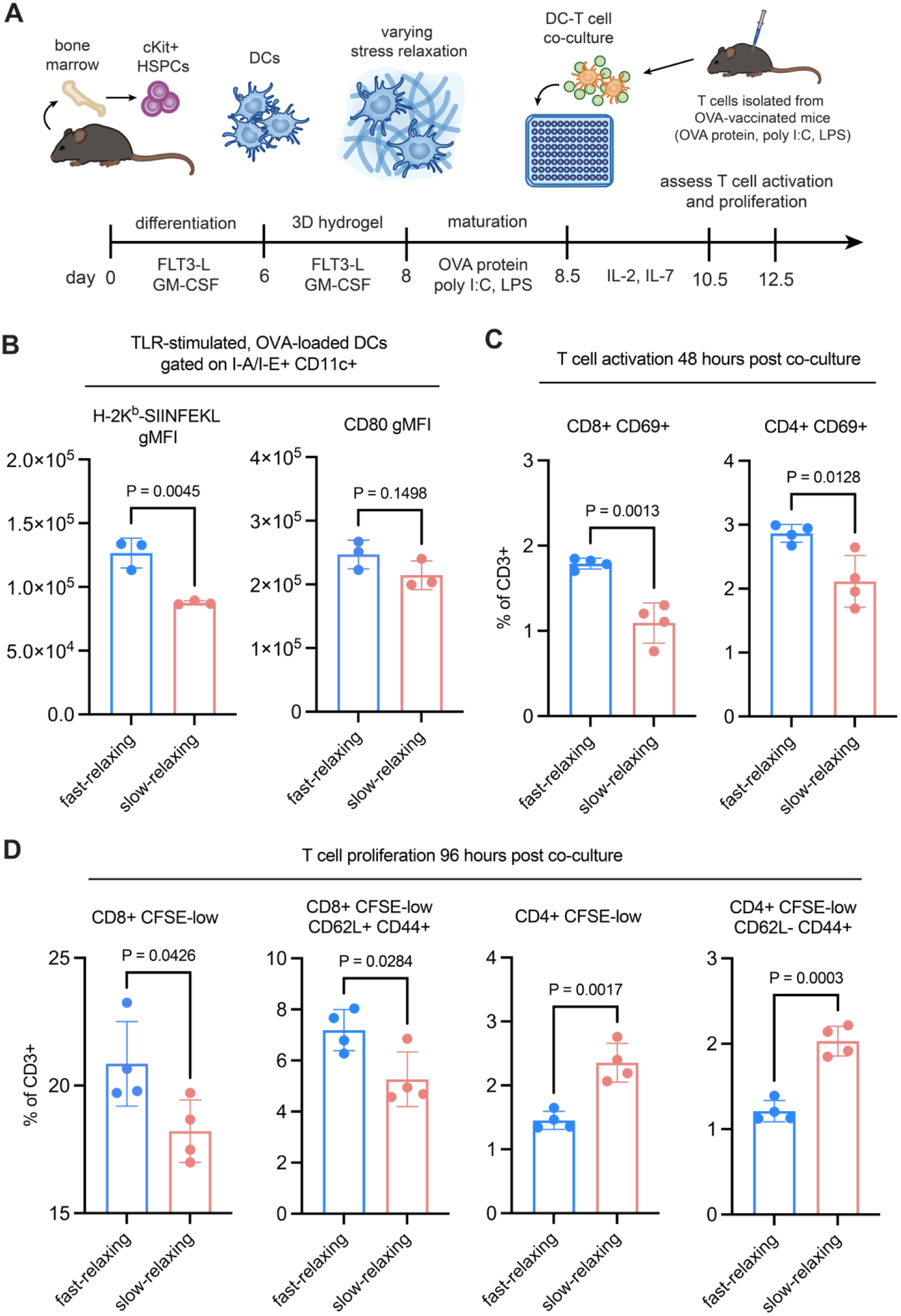
Matrix stress relaxation impacts antigen presentation and antigen-specific T cell activation of murine stem cell-derived DCs in vitro. **(A)** Experimental workflow for the in vitro assessment of mechanically primed murine DCs (mDCs). cKit+ progenitors were isolated from the long bones of C57BL/6 donor mice and differentiated into DCs. Mechanically educated mDCs were matured with TLR agonists, and loaded with OVA. OVA-reactive T cells were isolated from donor C57BL/6 mice vaccinated with OVA protein, poly I:C and LPS. OVA-presenting mature mDCs were co-cultured with OVA-reactive T cells for 48 and 96 hours to assess T cell activation and proliferation. **(B)** mDCs educated by the fast-relaxing matrix promoted presentation of the class I OVA peptide and CD80 post TLR activation. **(C)** Higher frequency of CD69+ activated T cells were induced by the fast-relaxing mDCs. **(D)** Fast-relaxing mDCs generated a CD8 bias T cell response with enhanced CD8+ proliferation and central memory response whereas slow-relaxing mDCs had higher CD4+ proliferation with short-lived effector response. Data in B are representative of n = 3 biological replicates from three independent hydrogel samples, and data in C and D are representative of n = 4 biological replicates from four independent hydrogel samples. Data are shown as mean ± SD, and P-values from two-tailed unpaired Student’s t-tests are indicated.

### Fast-relaxing DCs generate a CD8-biased antigen-specific T cell response in vitro

Activated mDCs loaded with OVA protein were co-cultured with OVA-reactive T cells to directly examine the T cell priming capacity of the mechanically educated DCs. OVA-reactive T cells were isolated from the spleen and draining lymph node (dLN) of OVA-vaccinated mice to include all possible epitopes of the OVA protein **(Fig. 4A)**. After 48 hours of co-culture, fast-relaxing mDCs induced more activated CD69+ T cells and Ki67+ CD8+ T cells **(Fig. 4C, S7B)**. Moreover, the CD4/CD8 ratio was lower in the fast-relaxing condition, suggesting a CD8-biased response that is consistent with the higher expression of MHC-I-OVA peptide complex in fast-relaxing mDCs **(Fig. S7A, 4B)**. After 96 hours of long-term co-culture, fast-relaxing mDCs promoted more CD8+ T cell proliferation, and the proliferated CD8+ T cells had a higher CD44+ CD62L+ central memory population **(Fig. 4D, S7E)**. Conversely, the slow-relaxing mDCs had higher CD4+ proliferation. These proliferating T cells were mostly the short-lived CD44+ CD62L-effector-like cells. These results suggest that fast-relaxing mDCs had the capacity to generate a potent memory CD8+ T cell response that is often associated with cancer remission and rapid response to reinfection^56,57^.

### Fast-relaxing DCs generate strong draining lymph node T cell responses through adoptive cell transfer in vivo

We next investigated whether our findings were translatable in vivo through adoptive transfer of mechanically educated DCs. After hydrogel cultures, mDCs were TLR-activated and loaded with OVA protein, followed by subcutaneous injection into the right flank of antigen-naïve mice **(Fig. 5A)**. After 24 hours, more I-A/I-E+ CD11c+ CD11b+ DCs were recruited to the draining lymph node (dLN) compared to the contralateral lymph node **(Fig. S8A)**. However, no differences in DC recruitment or T cell frequency were observed in the dLN across all DC groups **(Fig. 5B, S8B)**. Draining lymph nodes were harvested 10 days post-cell transfer to assess the T cell priming response from the injected DCs. Consistent with our in vitro result, fast-relaxing mDCs generated a CD8 T cell response with more class I tetramer-positive cells and increased CD8+ T cell frequency in the dLN **(Fig. 5C, S8C)**. Moreover, fast-relaxing mDCs induced higher frequencies of CD44+ CD62L- antigen-experienced effector-like T cells with no changes in the CD44+ CD62L+ central memory T cell frequency **(Fig. 5C)**. To test the T cell response to peptide restimulation, dLN cells were treated with class I (OVA_257-264_) and class II (OVA_323-339_) OVA peptides for 24 hours. Upon restimulation, dLN cells treated with the fast-relaxing mDCs had the highest frequencies of proliferating Ki67+ cells and activated CD69+ CD8+ T cells **(Fig. 5D, S8E)**. In addition, more IFN-γ secretion, primarily contributed by the differences in IFN-γ+ CD8+ cells, were observed in the fast-relaxing conditions **(Fig. 5E, S8F)**. The outcome of the in vivo studies demonstrated that mechanical education through the fast-relaxing 3D environment licensed DCs to prime dLN T cells with a CD8+ effector bias which were potent responders to restimulation.

**Figure 5.**
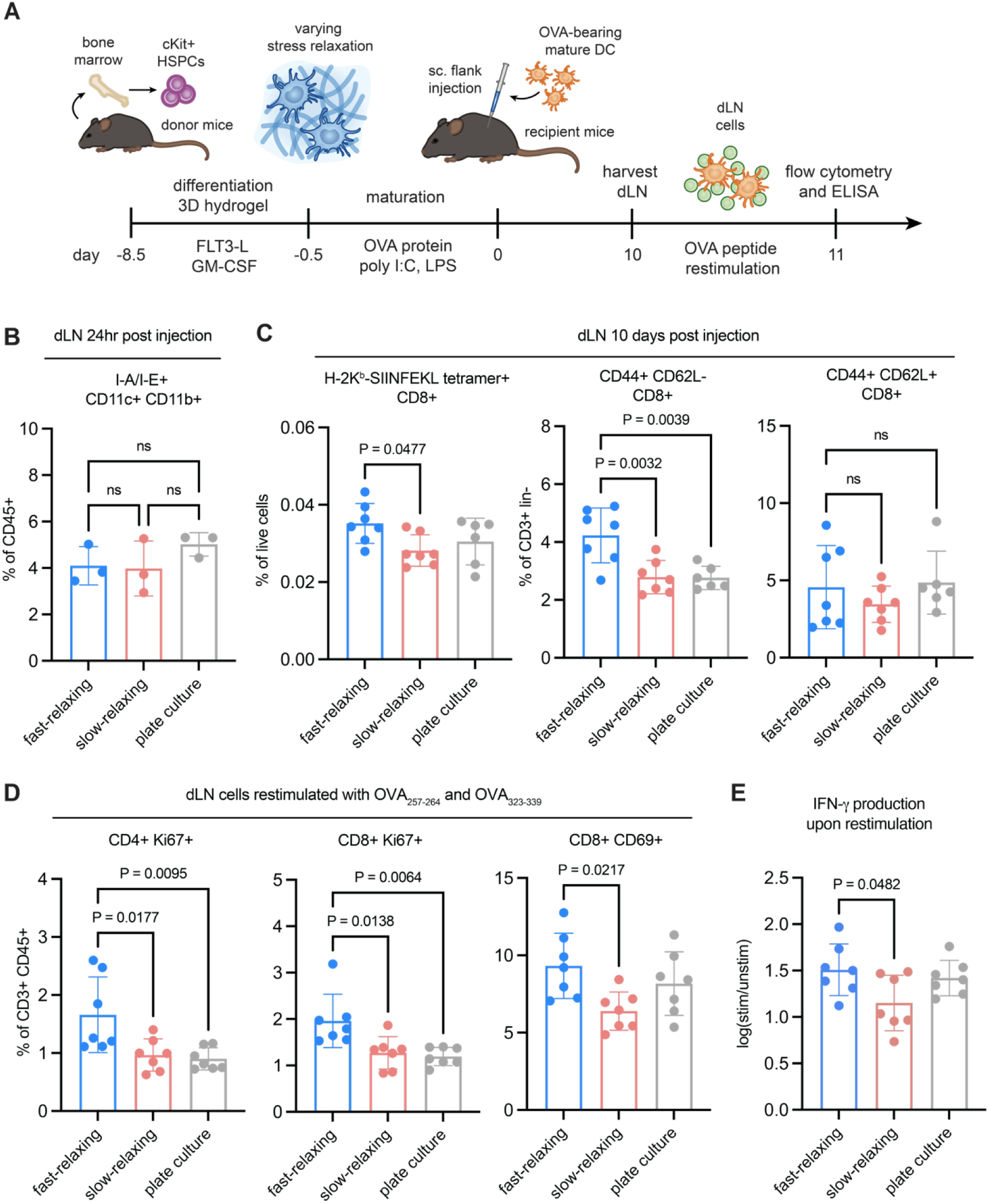
Mechanically educated DCs had differential in vivo T cell priming capacity through adoptive cell transfer. **(A)** Workflow for the in vivo assessment of mechanically primed murine DCs (mDCs). Mechanically educated DCs were matured with TLR agonist and loaded with OVA protein, followed by subcutaneous flank injection into C57BL/6 recipient mice. Draining lymph nodes (dLNs) were harvested at the study end point and restimulated with OVA peptides. **(B)** No statistical difference in MHC-II+ CD11b+ CD11c+ DC recruitment to the dLN was observed 24 hours after the injection. **(C)** Fast-relaxing mDCs generated more OVA tetramer-positive T cells in the dLNs after 10 days with higher frequency of CD8+ effector-like cells in the total T cell population. Lineage-negative (lin-) is defined by I-A/I-E- CD19- CD11c-. **(D)** Upon restimulation with both class I and II OVA peptides, dLN cells from the fast-relaxing DC condition showed the highest activated Ki67+ and CD69+ T cells. **(E)** IFN-γ production from dLN cells analyzed by ELISA. Expression was normalized by log (fold change) of stimulated to unstimulated cells. mDCs educated by the fast-relaxing matrix produced the most antigen-specific IFN-γ response. Data are shown as mean ± SD, and P-values from one-way ANOVA with Tukey’s post-hoc test are indicated. Data are representative of biological replicates shown as mean ± SD. P-values from one-way ANOVA with Tukey’s post-hoc test are indicated.

### Dendritic cells sense matrix stress relaxation through PI3K signaling and actin branching facilitated by cytokine signaling

Having established that mechanically educated DCs took on distinct cell fate and had differential T cell priming capacity, we aimed to examine cellular elements that contribute to mechanosensing. Morphological analysis of DCs in the hydrogels showed that the slow-relaxing hydrogel promoted a more rounded shape with increased cell volume and F-actin expression, indicating the role of actin cytoskeleton in responding to stress relaxation **(Fig. 6A)**. Since the frequency of HLA-DR+ CD11c+ CD80+ PD-L1+ cells were enriched in the slow-relaxing matrix, we tried to perturb this population through changes in cytokine signaling and pharmaceutical inhibition. Mechanotransduction pathways are often dependent on extracellular biochemical signaling^16,58^. To test whether cytokine signaling is essential for sensing matrix stress relaxation in DCs, IL-4 was removed from the culture media. GM-CSF was kept in the media to maintain the viability of DCs during hydrogel culture^35,59^. When DCs were only exposed to GM-CSF, the frequency of HLA-DR+ CD11c+ CD80+ PD-L1+ decreased in the slow-relaxing condition and had no statistical difference compared to the fast-relaxing hydrogel **(Fig. 6B)**. This observation demonstrated that the concerted action of IL-4 and GM-CSF signaling was required to drive cell state changes in DCs under different matrix stress relaxation.

**Figure 6.**
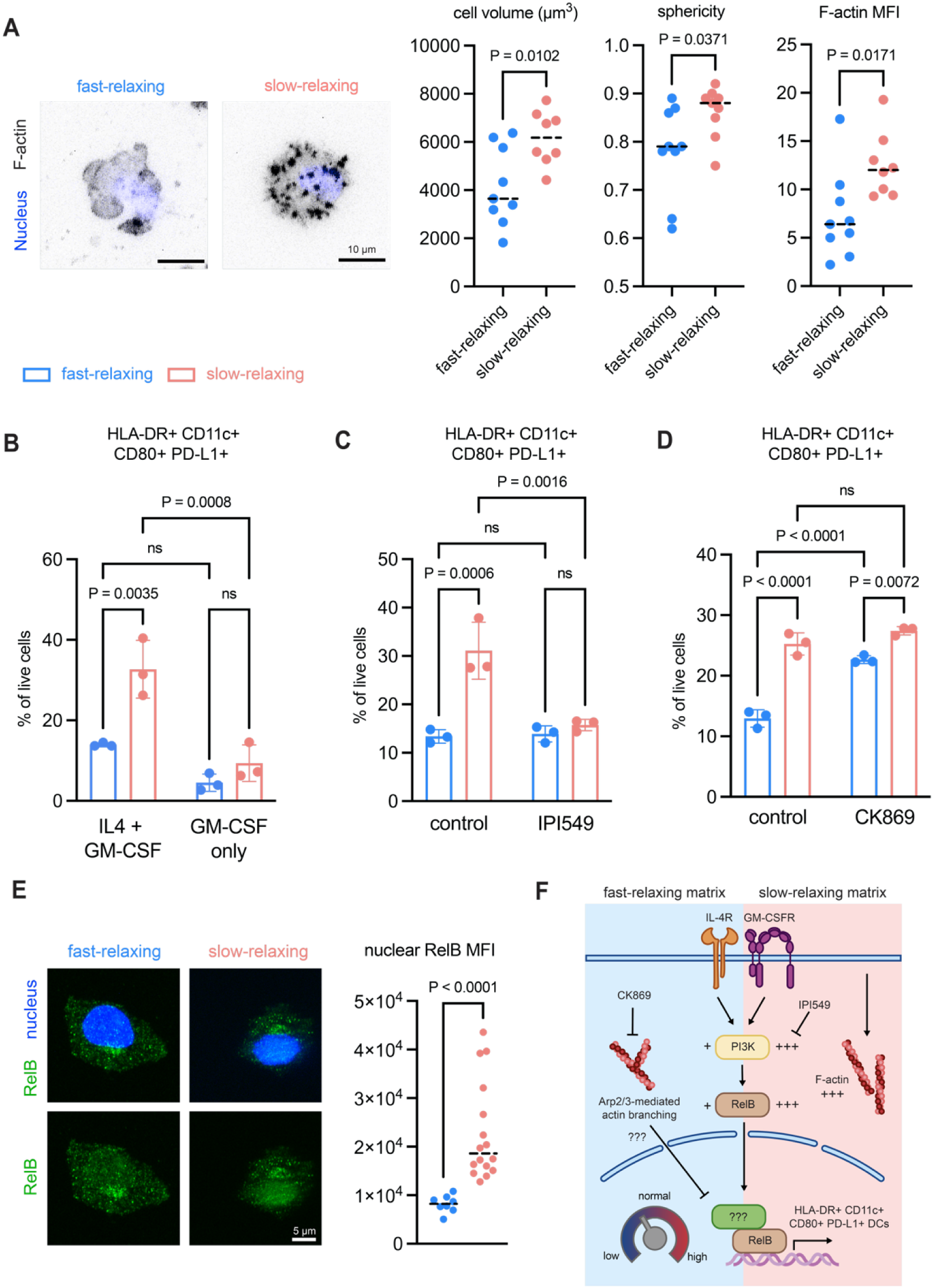
Mechanical education through matrix stress relaxation is dependent on PI3K signaling and F-actin branching. **(A)** Representative images of human DCs encapsulated in the fast-relaxing and slow-relaxing hydrogels. F-actin was stained with phalloidin (pseudocolored in black) and nuclei were stained with DAPI (blue). Quantifications of cell volume, sphericity, and F-actin mean fluorescent intensity (MFI) are represented as dot plots where each dot represents a single cell. DCs in the slow-relaxing matrix exhibited a larger, more rounded morphology with increased F-actin expression. **(B)** The slow-relaxing matrix promoted the frequency of HLA-DR+ CD11c+ CD80+ PD-L1+ DC state. This slow-relaxing state was dependent on the concerted cytokine signaling of IL-4 and GM-CSF. **(C)** The slow-relaxing state relied on PI3K signaling and was inhibited by IPI549 (γ isoform). **(D)** Inhibition of Arp2/3 complex using CK869 induced the slow-relaxing state in the fast-relaxing matrix. **(E)** Representative immunofluorescent images of DCs retrieved from fast-relaxing and slow-relaxing hydrogels. DCs from the slow-relaxing matrix exhibited an increased in nuclear RelB (green) localization compared to the fast-relaxing condition. Nuclei were stained with DAPI (blue). Quantification of nuclear RelB MFI is shown as dot plots where each dot represents a single cell. **(F)** Proposed schematic of mechanical regulation of DC cell state through matrix stress relaxation and cytokine signaling. P-values from two-tailed unpaired Student’s t-tests in A and E are indicated. Data in B, C, and D are representative of n = 3 biological replicates from independent hydrogel samples, and they are shown as mean ± SD. P-values from two-way ANOVA with Tukey’s post-hoc test in B, C, and D are indicated.

PI3K signaling pathway was upregulated in slow-relaxing DCs and was previously shown to contribute to viscoelasticity-sensing^33^ **(Fig. 3B)**. Its involvement is crucial for T cell priming and inflammation, but aberrant signaling leads to immune suppression and dysfunction^60–62^. Dendritic cell functions are often associated with the γ and δ isoform of PI3K, so we treated encapsulated DCs with pharmaceutical inhibitors specifically targeting these two isoforms^63^. PI3K inhibition regardless of the isoforms led to decreased frequency of HLA-DR+ CD11c+ CD80+ PD-L1+ DCs in the slow-relaxing matrix **(Fig. 6C, S9A)**. Interestingly, they became comparable to the fast-relaxing condition, and the inhibition had no effect in the fast-relaxing matrix. This result implied that reduced matrix stress relaxation activated PI3K signaling in DCs to alter their cell state. Because changes in F-actin were observed, we perturbed the actin cytoskeleton through pharmaceutical inhibition of the Rho-ROCK pathway using Y-27632. In addition, previous study showed that DCs alter their CCR7 expression through Arp2/3 complex-mediated sensing of their cell shape^64^, so Arp2/3 complex inhibitor CK869 was used to treat DCs in the hydrogel cultures. No significant differences were observed from Rho-ROCK inhibition, suggesting that actomyosin contractility did not play a role in stress relaxation-sensing in DCs **(Fig. S9B)**. On the contrary, inhibiting Arp2/3 complex showed an increase in HLA-DR+ CD11c+ CD80+ PD-L1+ frequency in the fast-relaxing hydrogels **(Fig. 6D)**. This surprising result indicated that disruption in actin branching contribute to the dysfunctional state of slow-relaxing DCs characterized by lower T cell priming capacity.

We analyzed cellular RelB expression using flow cytometry and RelB nuclear localization using immunocytochemistry to determine whether these matrix stress relaxation signaling cues have an impact on transcription factor expression. RelB is known as a member of the non-canonical NF-κB pathway which forms a heterodimer with p52 to mediate gene expression^65^. However, studies have shown that RelB integrates both the canonical and non-canonical pathways to regulate DC activation^66^. RelB acts as a rheostat of DC function where the degree of expression controls their tolerogenic phenotype and immune suppression^67–69^. By removing IL-4 from the media or treating with PI3Kγ inhibitor IPI-549, total RelB expression decreased in the slow-relaxing matrix, which corresponds to the reduction of HLA-DR+ CD11c+ CD80+ PD-L1+ population **(Fig. S9C, S9D)**. However, treatment with CK869 to inhibit Arp2/3 complex had no effect on total RelB expression **(Fig. S9E)**. Nuclear expression of RelB was higher in the slow-relaxing DCs, indicating a heightened RelB-mediated transcription program in response to changes in matrix stress relaxation **(Fig. 6E)**. Surprisingly, as opposed to the increased HLA-DR+ CD11c+ CD80+ PD-L1+ population observed with Arp2/3 complex inhibition in fast-relaxing DCs, CK869 treatment did not increase nuclear RelB expression in fast-relaxing DCs, and instead led to a reduction in nuclear expression **(Fig. S9F)**. We hypothesized that the distinct expression of RelB observed in similar phenotype can be attributed to the rheostat analogy where both over-expression and under-expression of RelB leads to a dysfunctional state of DCs^67–69^. Collectively, these data suggest that while both PI3K signaling and actin branching serve as sensors of matrix stress relaxation in DCs, they regulate DC fate through distinct mechanisms **(Fig. 6F)**.

## Conclusion and Outlook

This study demonstrates that the 3D mechanical environment, particularly matrix stress relaxation, acts as a factor contributing to the heterogeneity of dendritic cells and their ability to mount T cell responses. Using a 3D hydrogel system with tunable stress relaxation, we showed that DCs cultured in the fast-relaxing matrix exhibited superior antigen presentation and CD8 T cell priming both in vitro and in vivo. Conversely, DCs from the slow-relaxing matrix had a hyper-inflammatory phenotype with reduced capacity to present antigen, potentially caused by lipid accumulation and lack of regulatory mechanisms. Notably, the differences in DC states persisted with TLR activation even after the removal of mechanical cues and TLR activation. These data suggested that mechanical cues license DC states that can lead to differential outcomes to CD8-mediated inflammatory, anti-viral, or anti-tumor response. We further established that DCs sensed matrix stress relaxation through PI3K signal transduction and actin branching using pharmaceutical inhibition, and changes in DC state were dependent on concerted cytokine signaling. Moreover, RelB activity was shown to be correlated with the dysfunctional state observed in the slow-relaxing matrix, where both over-expression and under-expression of RelB in the nucleus lead to the same phenotype. Future studies can focus on the regulation of RelB-mediated transcription to gain insights into the relationship between matrix stress relaxation, signaling pathways, actin cytoskeleton, and transcription factor regulation. Targeting dendritic cells has become a popular immunotherapy approach in treating cancer and autoimmune diseases. However, the lack of understanding towards how DCs acquire heterogenous states hinders the development of these therapies. This study proposes a new mechanism towards DC heterogeneity through mechanical regulation of changes in matrix stress relaxation, potentially pointing out novel methods to generate dendritic cell-based cellular therapy.

## Methods

### Cell culture

De-identified human G-CSF-mobilized CD34+ HSPCs were purchased from Fred Hutchinson Cancer Center. HSPCs were cultured at 100k/ml in StemSpan^TM^ SFEM II (StemCell Technologies) supplemented with recombinant human IL3 (20 ng/mL, Peprotech), recombinant human stem cell factor (100 ng/mL, Peprotech), recombinant human FLT3 ligand (100 ng/mL, Peprotech), recombinant human thrombopoietin (50 ng/ml, Peprotech), and small molecule UM729 (700 nM, Stem Cell 15 Technologies) in U-bottom 96-well plates for 5 days per expansion passage. Half media changes were performed with freshly supplemented cytokines every 3 days.

For differentiation, expanded human HSPCs were seeded at 100k/ml in Iscove’s modified Dulbecco’s medium (IMDM Modified, Gibco) with 10% heat-inactivated fetal bovine serum (Gibco), non-essential amino acid (Gibco), and Antibiotic-Antimycotic (Gibco) supplemented with human recombinant FLT3-L (100 ng/ml, Peprotech) and human recombinant GM-CSF (100ng/ml, Peprotech) in U-bottom 96-well plates for 7 days. Half media changes were performed with freshly supplemented cytokines every 3 days. During encapsulation in the hydrogels, DCs were cultured in the IMDM media described above supplemented with recombinant human IL-4 (50 ng/mL, Peprotech) and GM-CSF (50 ng/mL, Peprotech) for 5 days. A complete media change was performed at day 3.

To generate murine DCs, cells were isolated from the long bones of female C57BL/6 mice (4 weeks old) following previously published protocols, and cKit+ progenitor cells were isolated using the EasySep™ Mouse CD117 (cKIT) Positive Selection Kit (StemCell Technologies). To differentiate cKit+ progenitor cells into DCs, they were seeded at 200k/ml in the IMDM media described above supplemented with murine recombinant FLT-3L (50 ng/ml, Peprotech) and murine recombinant GM-CSF (50 ng/ml, Peprotech) for 6 days. At day 3 of culture, same volume of media with freshly supplemented cytokines was added. During encapsulation in the hydrogels, mDCs were cultured in media with the same formulation as differentiation. To activate mDCs, they were seeded at 1M/ml in IMDM media supplemented with poly I:C (HMW) (10 µg/ml, InvivoGen), ultrapure LPS (LPS-EB) (100 ng/ml, InvivoGen), and OVA protein (EndoFit™ OVA protein) (100 µg/ml, InvivoGen) for 15 hours.

All cell cultures were maintained at 37°C in a humidified chamber with 5% CO2. Mycoplasma was routinely screened using MycoAlert. Cells counts were performed using a hemocytometer.

### Materials synthesis and preparation

Click-modified alginate was prepared as previously published. In brief, very low molecular weight ultra-pure sodium alginate (Provona UP VLVG, NovaMatrix) was covalently coupled with either (4-(1,2,4,5-tetrazin-3-yl)phenyl)methanamine hydrochloric acid (Tz, KareBay Biochem) or 5-(aminomethyl)bicyclo[2.2.1]hept-2-ene (Nb, norbornene methanamine, TCI America). The reaction product was filtered (0.22 µm), centrifuged, and lyophilized for long-term storage. All chemicals were purchased from Sigma-Aldrich. The interpenetrative network was prepared as previously published. In brief, bovine telo-collagen type I (BIOMATRIX) was neutralized using 1 M sodium hydroxide, and the final concentration of type I collagen was 4 mg/ml. Then, 10x Hanks’ balanced salt solution (HBSS, Gibco) and 1 M N-2-hydroxyethylpiperazine-N-2-ethane sulfonic acid (HEPES, Gibco) were added to the solution. Calcium carbonate slurry from precipitated calcium carbonate (Multifex-MM, Specialty Minerals) was prepared using sonication and added to the mixture. Then, unmodified or modified alginate was added based on the desired properties to the final weight concentration of 1 wt%. To initiate gelation, glucono-δ-lactone dissolved in HBSS/HEPES (EMD Millipore) was added as the very last step, and gels could be casted for desired usage. For cell encapsulation experiments, cells were resuspended with the 1x HBSS/HEPES buffer and added into the neutralized collagen solution.

### Oscillatory rheological characterization of interpenetrative hydrogel networks

Rheological characterization of the interpenetrative network was performed on a stress-controlled rheometer (HR-30, TA Instrument) using a 20 mm parallel plate. Gel solution was casted on sand-blasted Peltier plate at 37°C with a solvent trap to prevent dehydration. Time sweep was performed under 2% strain and 0.1Hz to measure plateau materials properties such as initial storage modulus and tan(delta). After complete gelation, a shear stress relaxation test at 10% strain applied in 5 seconds was performed to measure the decay of stress overtime under a constant strain.

### Cell retrieval from the interpenetrative networks

Hydrogels were digested with 900 U ml−1 collagenase I (Worthington) and 34 U ml−1 of alginate lyase (Sigma-Aldrich) in DBPS supplemented with calcium, magnesium (Gibco), and 0.5% BSA (Sigma-Aldrich) at 37 °C for 20 min, followed by addition of another 900 U ml−1 collagenase I and incubation at 37 °C for an hour. After incubation, cells were washed in DPBS supplemented with 2 mM EDTA (Sigma-Aldrich) and 0.5% BSA (Sigma-Aldrich) by centrifuging at 350g for 5 minutes at 4°C twice. A final wash was done in DPBS with centrifuging at 350g for 5 minutes at 4°C.

### Flow cytometry analysis

Prior to antibody staining, cells were stained with Live/Dead Fixable blue dye (Thermo) for 20 minutes at 4°C followed by blocking with TruStain human or mouse FcX (Fc Receptor Blocking Solution, Biolegend) for 10 minutes at 4°C. Then, cells were incubated with a mix of antibodies for 20 minutes at 4°C. A table of flow antibodies and dilutions used in this work is included in the Supplementary Materials. Cells were fixed at 2% Paraformaldehyde (Electron Microscopy Sciences) in DPBS for 10 minutes at RT and resuspended in FACS buffer (Invitrogen) at 4°C for short-term storage. For intracellular staining, fixed cells were permeabilized with 0.2% Triton-X and 0.5% BSA in DPBS for 10 minutes at 4°C. Then, antibodies for intracellular target were incubated for 20 minutes at 4°C, and cells were resuspended in FACS buffer for short-term storage. Stained cells were analyzed using the Cytek Aurora 5-laser Spectral Flow Analyzer (Flow Cytometry Core at the Children’s Hospital of Philadelphia) and FCS Express (De Novo Software). Unstained control and single-color control using UltraComp compensation beads (Thermo) were included in the analysis, and gating was performed using the unstained control. Gating strategies is included in the Supplementary Materials.

### 3D cell migration assay

Encapsulated cells were retrieved as described above and resuspended in the IMDM media. Isolated cells were resuspended in neutralized type I bovine telo collagen solution (1mg/ml in HBSS/HEPES buffer, BIOMATRIX) at 1M cells/ml. The gel construct rested at 37°C in the humidified incubator overnight. Prior to imaging, cells were stained with calcein AM (Thermo). Live cell imaging was performed using time lapse with z-stack on a confocal microscope (Leica Stellaris, Penn Vet Imaging Core). Z-stacks were performed every 5 minutes and each gel was imaged for 40 minutes in total. Cell track analysis was done using the TrackMate plugin of Fiji in the green channel. Spider plots were generated using the track data from TrackMate and plotted in Graphpad Prism.

### Phagocytosis assay

DCs retrieved from the hydrogels were resuspended in the IMDM media at 100k/mL and replated on MatTek glass bottom dishes with 10mm microwell (Fisher) for 1 hour. Then, pHrodo Red E Coli. Bioparticles Conjugate beads (Fisher) were added to the plates at 100 µg/ml, and DCs were exposed for 2 hours at 37°C. The reaction was quenched by replacing the media with DPBS. CellMask Green (Fisher) was used to stain for cell boundary for imaging analysis, and cells were gently fixed with 2% PFA (Electron Microscopy Sciences) in DPBS after staining. Images were taken on EVOS M000 fluorescent microscope (ThermoFisher) at 20x and analysis was done through ImageJ. In brief, signals in the green channel was used to generate cell mask that indicate individual cell boundary through particle analysis. Overlapping cells were excluded based on the cell area. Then, fluorescent signals from the red channel was measured using the masks. The red signals were blank-subtracted and normalized by the area of each mask.

### TLR activation and ELISA assays

DCs retrieved from hydrogels were resuspended with IMDM media supplemented with 10 µg/ml poly I:C (TLR3, Invivogen), 1 µg/ml ultrapure LPS (TLR4, Invivogen), or 2.5 mM CpG ODN M362 (TLR9, Invivogen), and replated in U-bottom 96-well plates at 150k/ml for 24 hours. Then, conditioned media was collected for ELISA analysis. Activated cells were processed for flow cytometry described above. Media samples were stored at -80°C for short-term storage. Diluted samples were used to perform ELISA assays following manufacturer’s protocol. All ELISA kits were purchased from Biolegend.

### Mixed lymphocyte assay

DCs retrieved from hydrogels were resuspended with IMDM media supplemented with 10 µg/ml poly I:C (Invivogen) and replated in U-bottom 96-well plates at 50k/ml for 24 hours. Allogeneic CD3+ T cells purchased from the Human Immunology Core at the University of Pennsylvania were stained with CFSE cell tracker kit (0.5 µM, Biolegend) following the manufacturer’s protocol. CFSE-stained T cells were resuspended in RPMI 1640 medium (Gibco) with 10% heat-inactivated detal bovine serum (Gibco), HEPES buffer (20mM, Gibco), GlutaMAX^TM^ supplement (Gibco), and Antibiotic-Antimycotic (Gibco), supplemented with recombinant human IL-2 (20 ng/ml, Peprotech). The two cell types were added to each well in a ratio of 1:10 (DC:T) with 500k/ml T cells for the 4-day co-culture. A half-media change was performed at day 2 and cells from each well were processed at day 4 for flow cytometry described above.

### RNA isolation and bulk RNA sequencing analysis

DCs retrieved from hydrogels without TLR activation are processed with the RNeasy Micro Kit (Qiagen) to obtain RNA. RNA samples were stored at -80°C for short-term storage. Library preparation, sequencing, and alignment were performed by Novogene along with quality control for RNA and sequencing. All sequencing analysis was done using R studio. Principle component analysis was performed using the EdgeR package and prcomp function, and the plot was generated using ggplot. Differentially expressed genes were analyzed using the EdgeR and limma package. P-values are FDR-adjusted using Benjamini-Yekutieli method. DEGs are filtered based on adjusted P values < 0.05 and log fold change > 1. Gene Ontology term enrichment analysis was done with the clusterProfiler package. P values were FDR-adjusted using Benjamini-Hochberg method and GO terms were filtered with p value lower than 0.05 and q value lower than 0.2. Heatmaps of gene sets were generated using the pheatmap function with row normalization. TCGA datasets were extracted using the TCGAbiolinks package. The cohort included all available RNA count data from BRCA, COAD, HNSC, LUSC, and SKCM cancer subtype. Enrichment scores of each gene set were calculated using ssGSEA in the GSVA pacakage. All gene sets were calculated in the same execution. Gene sets include upregulated DEGs, downregulated DEGs, CTL (PRF1, GZMA, GZMB, GNLY, IFNG, CD8A, CD8B, KLRK1, FASL) and Th1 (IFNG, TBX21, STAT1, STAT4, IL12RB1, CXCR3, CCR5, IFNGR1, IL2). The spearman correlation between gene sets in each sample was obtained using the cor.test function in R. Trendline in the scatter plot was added using abline and lm functions. Small P-values that were beyond computational limit were plotted as 10^-320^. CIBERSORT analysis was performed on the CIBERSORTx website. Published dendritic cell single cell sequencing data (GSE94820) was uploaded as a signature matrix, and bulk RNA sequencing data from the mechanically primed DCs was uploaded as a mixture file. Deconvolution was run using S-mode batch-correction in the relative mode (forcing the sum of fraction to be 100) with 100 permutations. Deconvolution results were ensured to have P-value < 0.05.

### Generation of OVA-T cells and antigen-specific DC-T cell co-culture

To generate murine OVA-reactive T cells, a cocktail of OVA protein (EndoFit™ OVA protein) (50 µg, InvivoGen), poly I:C (HMW) (25 µg, InvivoGen), and ultrapure LPS (LPS-EB) (10 µg, InvivoGen) was subcutaneously injected in the right flank of naïve female C57BL/6 mice (6 weeks old, Taconic Biosciences) on day 0 and 10. The draining lymph nodes and spleens were dissected on day 14, and T cells were isolated using EasySep™ Mouse T Cell Isolation Kit (StemCell Technologies). Isolated T cells were stained with CFSE cell tracker kit (0.5 µM, Biolegend) following the manufacturer’s protocol. 250k of CFSE-labeled T cells were co-cultured with 50k of OVA-presenting mature DCs (described above) in the complete RPMI media. Negative control contained no DCs and positive control had plate coated anti-CD3 and anti-CD28 (2ug/ml, 4°C overnight). Cells were retrieved for flow cytometry analysis after 48 and 96 hours to assess T cell activation and proliferation.

### In vivo adoptive cell transfer studies

500k activated, OVA-loaded murine dendritic cells (described above) were resuspended in 50 µL DPBS (gibco) and subcutaneously injected into the right flank of female C57BL/6 mice (8 weeks old). After 24 hours, the inguinal lymph nodes (draining and contralateral) were harvested (n = 3) to measure dendritic cell recruitment in the draining lymph node, and single-cell suspension of lymph node cells was made in FACS buffer (Invitrogen). After 10 days, the draining lymph nodes were harvested (n = 7) and made into single-cell suspension. Tetramer staining was done using the biotoin-H2K(b)-OVA monomer (Flex-T™ Biotin H-2 K(b) OVA Monomer (SIINFEKL), Biolegend) following the manufacturer’s protocol. Briefly, biotinylated MHC-peptide monomers were conjugated with either APC-streptavidin (Biolegend) or FITC-streptavidin (Biolegend) in a 4:1 ratio, followed by quenching the reaction with 40 µM of free d-Biotin. dLN cells were stained Live/Dead Fixable blue dye (Thermo) for 20 minutes at 4°C followed by blocking with TruStain mouse FcX (Fc Receptor Blocking Solution, Biolegend) for 10 minutes at 4°C. Tetramers of both colors were added at a dilution of 1:25 (FITC) and 1:50 (APC) for 20 minutes at 4°C in the dark. Then, other surface markers were added into the tetramer staining solution and incubated for another 20 minutes. Tetramer-positive cells were gated on double-positive cells in the FITC and APC channel to reduce non-specific signals. To restimulate with antigens, 500k draining lymph node cells were replated in RPMI media described above with 5 µg/ml of OVA 257-264 SINFEKL (VWR) and OVA 323-339 (VWR) for 24 hours at 37°C. Conditioned media was collected for IFN-γ ELISA (Biolegend) and cells were stained for flow cytometry.

### Immunofluorescent staining, confocal imaging, and image analysis

DCs encapsulated in the hydrogels were fixed in 4% PFA (Electron Microscopy Sciences) in DPBS at RT for 20 minutes followed by staining with Phalloidin 488 (1:1000, Abcam) and DAPI (1:2000, Abcam) at 4°C for 30 minutes on a rocker. Stained cells in the hydrogels were visualized under a confocal microscope (Leica Stellaris, Penn Vet Imaging Core). The morphology of the encapsulated cells and the actin intensity was analyzed using the IMARIS software (Penn Vet Imaging Core). For the RelB immunostaining, cells were first retrieved from the hydrogels using the same method described above. Single cell suspension in DPBS was collected in a V-bottom 96-well plate. Cells were fixed in 2% PFA in DPBS at RT for 20 minutes and permeabilized with 0.1% Triton-X (Sigma-Aldrich) in DPBS at RT for 15 minutes. To reduce non-specific signals, cells were blocked with 1% powdered bovine serum albumin (Sigma-Aldrich) and 0.5% normal goat serum (Abcam) in 0.05% Tween (Sigma-Aldrich) solution (staining buffer) for 1 hour at RT. Monoclonal rabbit anti-RelB (D7D7W) (Cell Signaling Technology) (1:400 in staining buffer) was added for 24 hours at 4°C on the rocker. After primary antibody staining, cells were washed with 0.05% Tween in DPBS for 3 five-minute intervals on the rocker. Alexa Fluor 594 Goat anti-Rabbit IgG (H+L) (ThermoFisher) (1:500 in staining buffer) was incubated for 24 hours at 4°C on the rocker followed by three washing steps. Cells were counterstained and mounted with ProLong Gold Antifade Mountant with DNA Stains (Invitrogen). Single-cell suspension was placed on a microscope slide covered with a circular coverslip for confocal imaging (Leica Stellaris, Penn Vet Imaging Core). Nuclear RelB intensity was measured using the IMARIS software.

### Cytokine and inhibition studies on mechanically primed DCs

For the cytokine study on IL-4 and GM-CSF, the control group was treated with the IMDM media supplemented with 50 ng/ml of IL-4 and GM-CSF describe above. The GM-CSF only group was treated with the IMDM media supplemented with 50 ng/ml of GM-CSF only. For the inhibition studies, 1 µM of IPI-549 (VWR International), 1 µM of idelalisib (Fisher), 1 µM of CK869 (Sigma), or 10 µM of Y-27632 (Selleckchem) was added to the IMDM media supplemented with 50 ng/ml of IL-4 and GM-CSF. The control groups were treated with the cytokine-supplemented IMDM media with 1 µM of DMSO (Thomas Scientific). Freshly added cytokines and inhibitors were added at day 3 of culture during the complete media change. Cells were retrieved from the hydrogels using the methods described above for flow cytometry staining. RelB intracellular staining for flow cytometry was performed using the methods above. After permeabilization, cells were first stained with the Monoclonal rabbit anti-RelB (D7D7W) (Cell Signaling Technology) (1:400 in staining buffer) for 20 minutes at 4°C, then washed twice by centrifuging at 350g. Then, cells were stained with the Alexa Fluor 594 Goat anti-Rabbit IgG (H+L) (ThermoFisher) (1:500 in staining buffer) for another 20 minutes at 4°C, followed by two washes.

## Author Contributions

Y.C. contributed to conceptualization, data curation, investigation, formal analysis, methodology, visualization, and writing. A.S.B., F.T-J., K.Z., N.T., S.Z., and H.M. contributed to investigation, formal analysis, and methodology. K.H.V. contributed to conceptualization, methodology, funding acquisition, supervision, visualization, writing, and resources.

## Data Availability

All data will be available from a publicly posted Dryad repository upon acceptance of the article at a DOI to be provided.

## Acknowledgement

This work was supported in part by the National Institute of Dental and Craniofacial Research of the National Institutes of Health (NIH) under Award Numbers R00DE030084 and L70DE035343 (KV), the National Institute of General Medical Sciences of the NIH under Award Number R35GM157079 (KV), the Research Scholar Grant, RSG-23-1152051-01-MM, from the American Cancer Society, https://doi.org/10.53354/ACS.RSG-23-1152051-01-MM.pc.gr.175462 (KV), and the Graduate Research Fellowship from the National Science Foundation, DGE-2236662 (NT, AB). HM was supported by a training grant to the Center for Innovation & Precision Dentistry (CiPD) (R90DE031532). Confocal images were taken using the Leica Stellaris 8 FALCON from the Penn Vet Imaging Core supported by NIH S10 OD032305-01A1. The authors thank Emily Cento, Zhilin Chen, Max A. Eldabbas, and Emileigh Maddox of the Human Immunology Core and the Division of Transfusion Medicine and Therapeutic Pathology at the Perelman School of Medicine at the University of Pennsylvania for providing de-identified CD3+ T cells that were purified from healthy donor apheresis using StemCell RosetteSep™ kits. The HIC is supported in part by NIH P30 AI045008 and P30 CA016520. HIC RRID: SCR_022380. The authors thank Ling Qin and Yanhua Lan for providing murine long bones for cKit+ cell isolation.

