## Supplementary Information for "Mechanical licensing of functional dendritic cell states for enhanced T cell priming"

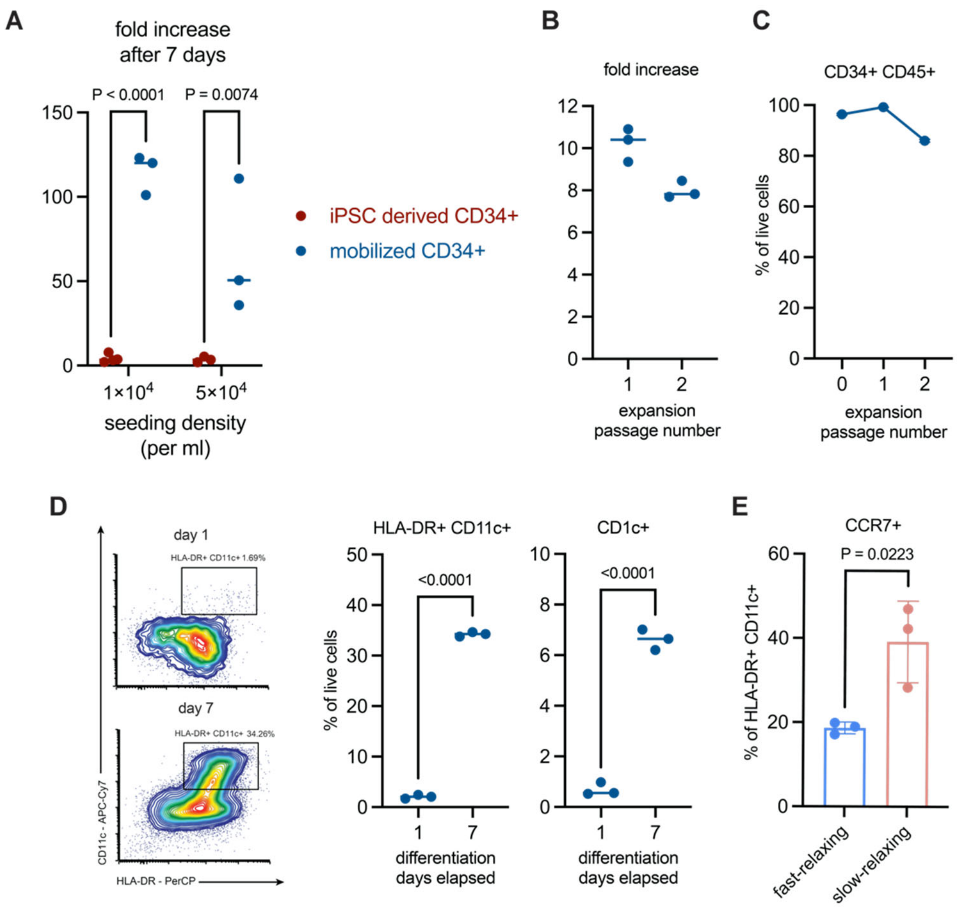


**Supplementary Figure S1 Related to Fig. 1 . (A)** Expansion of CD34+ HSPCs from two different sources. G-CSF mobilized CD34+ HSPCs had more expansion at varying seeding densities. This initial experiment was done with the addition of IL-6. **(B)** Fold increase of mobilized HSPCs in the expansion cytokine cocktail at each five-day passages. **(C)** Over 80% of CD34+ population was maintained after two expansion passages. **(D)** Representative flow cytometry plot of HLA-DR and CD11c expression on HSPCs in the differentiation cytokine cocktail. After 7 days of differentiation, the frequency of HLA-DR+ CD11c+ DCs increased with some committing to the CD1c+ lineage. **(E)** After 5 days of 3D hydrogel culture, HLA-DR+ CD11c+ DCs in the slow-relaxing matrix exhibited higher frequency of CCR7+ population. Data in A, B, C, D, and F indicate n = 3 biological replicates from the same human donor. Data in E indicate n = 3 individual hydrogel samples. Data are shown as mean ± SD, and P-values from two-tailed unpaired Student’s t-tests are indicated.

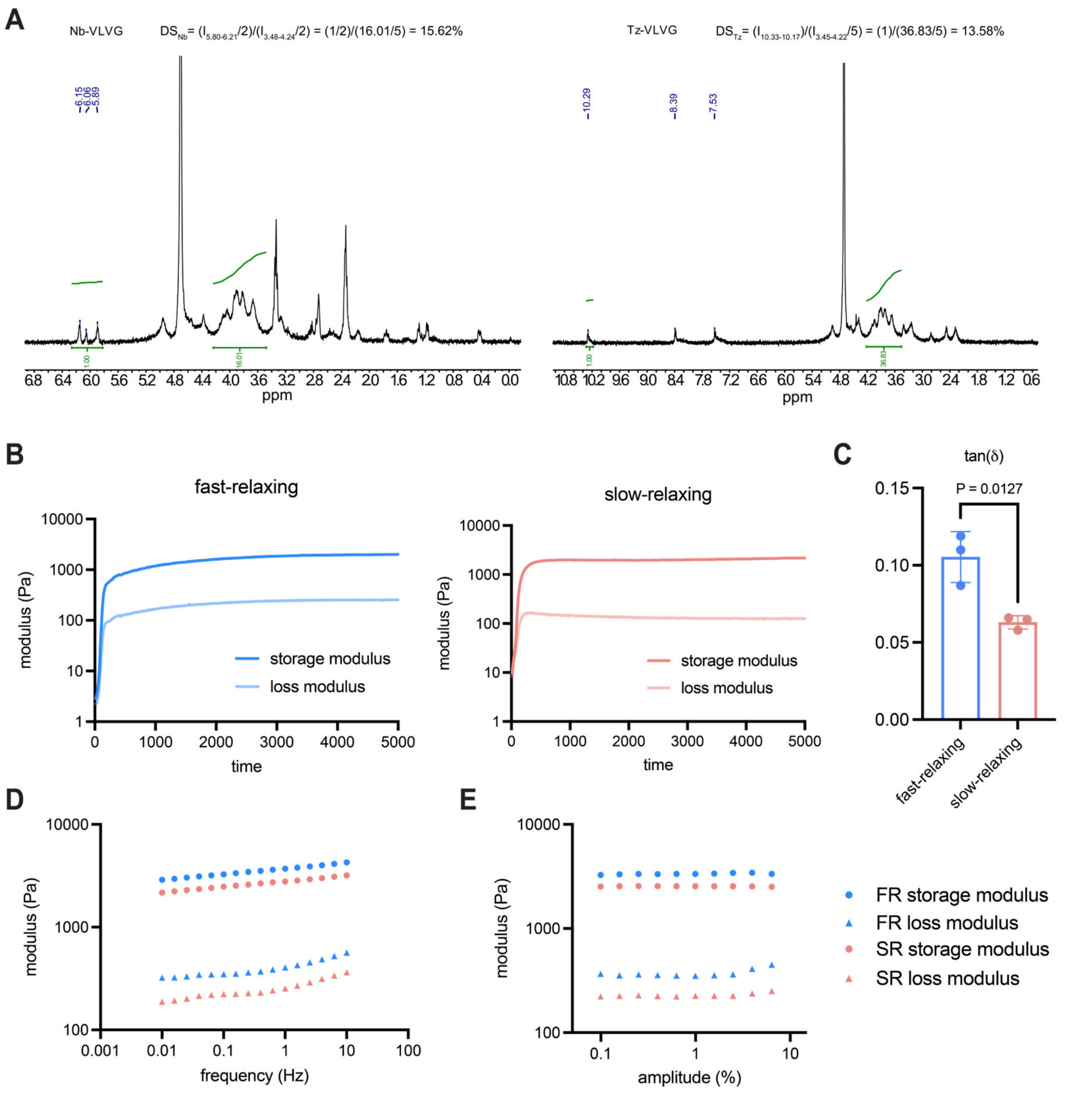


**Supplementary Figure S2 Related to Fig. 1 . (A)** NMR spectrum of alginate modified by norbornene (Nb, left) and tetrazine (Tz, right). The substitution rate was 15.62% for Nb and 13.58% for Tz. **(B)** Time sweep of fast-relaxing and slow-relaxing hydrogels. Both hydrogels reached stable modulus after an hour. **(C)** Slow-relaxing hydrogels had lower tan(delta) compared to fast-relaxing hydrogels, which is consistent with their stress relaxation. **(D)** Frequency sweep of both hydrogels from 0.01 to 10 Hz. Blue indicates fast-relaxing hydrogel and red indicates slow-relaxing hydrogel. Circular symbols indicate storage modulus whereas triangular symbols indicate loss modulus.

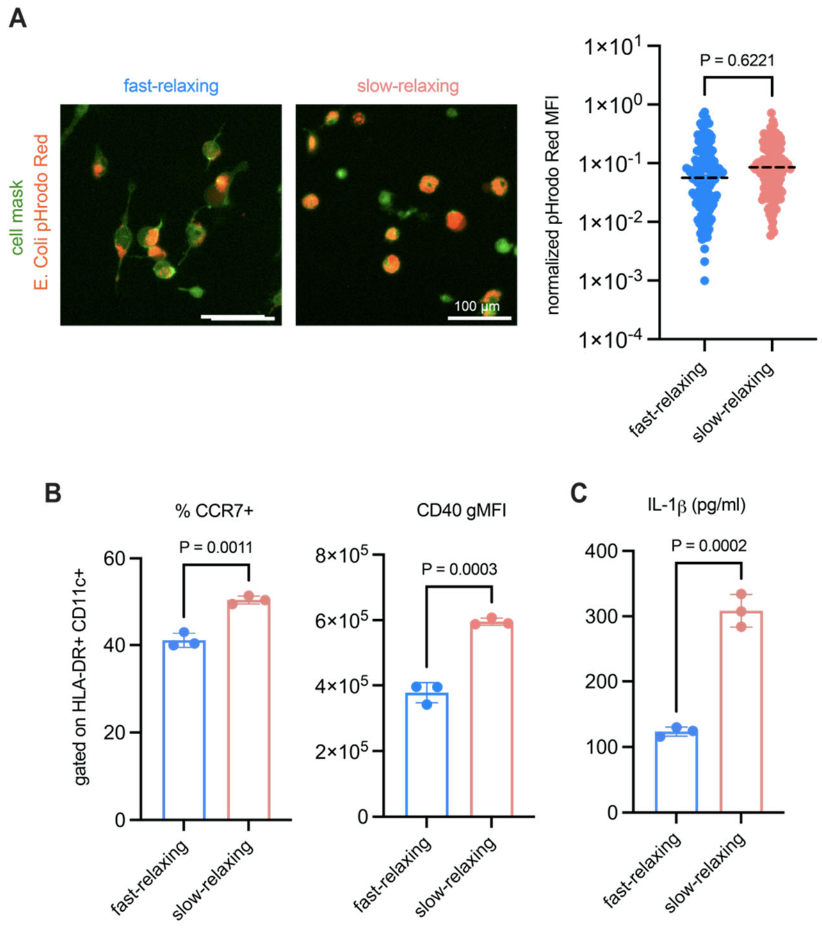


**Supplementary Figure S3 Related to Fig. 2. (A)** Representative images of mechanically educated DCs during phagocytosis. DCs were retrieved from hydrogels and reseeded in tissue culture plastic. Replated DCs were pulsed with pHrodo Red beads that fluoresced at low pH in the lysosome. The corresponding dot plot quantified the mean fluorescent intensity (MFI) of pHrodo beads within each cell, and no statistical difference in phagocytic activity were observed. **(B)** Post TLR3 activation, DCs from the slow-relaxing matrix had higher frequency of CCR7+ population and CD40 expression. **(C)** Slow-relaxing DCs promoted secretion of pro-inflammatory IL-1β secretion, consistent with the trend of IL-6 and TNF-α. Data in B and C are representative of n = 3 biological replicates from different hydrogels using the same human donor. Data in bar graphs are shown as mean ± SD, and P-values from two-tailed unpaired Student's t-tests are indicated.

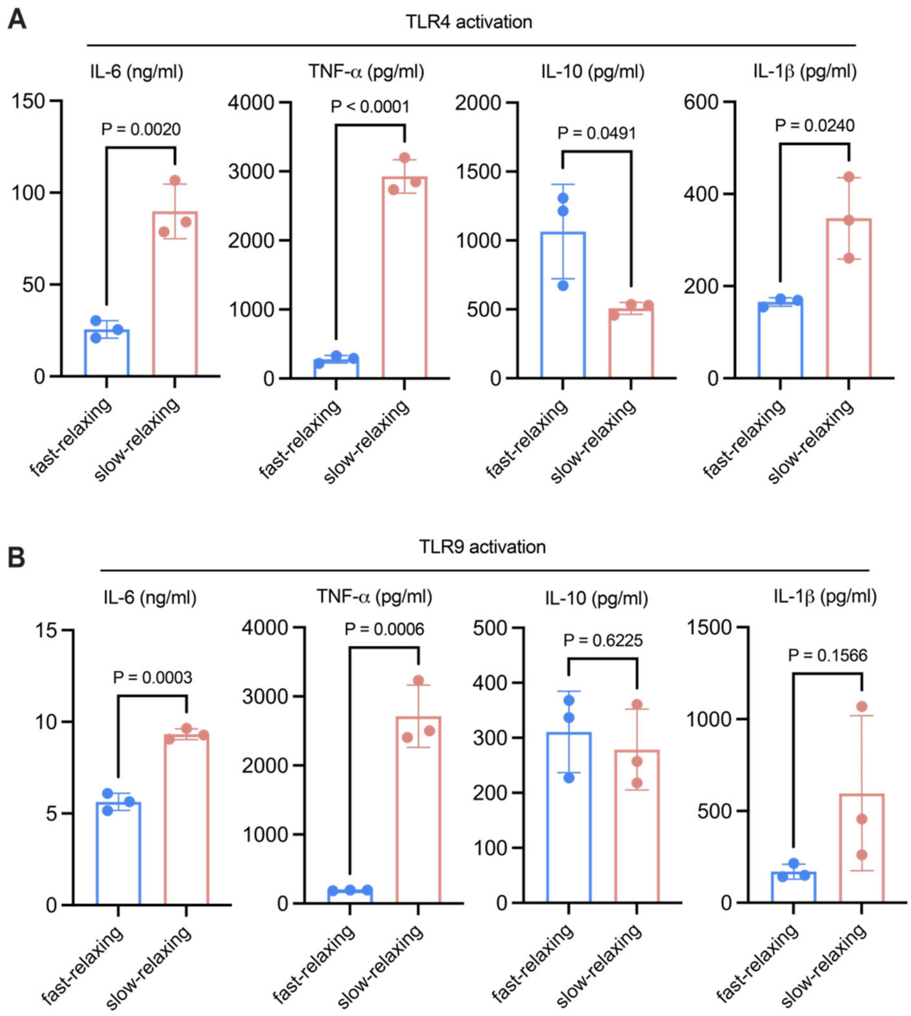


**Supplementary Figure S4 Related to Fig. 2.** Cytokine secretion upon **(A)** TLR4 and **(B)** TLR9 activation. Trends were mostly consistent as TLR3 activation where DCs educated by the slow-relaxing matrix promoted a pro-inflammatory cytokine bias. Data are representative of n = 3 biological replicates from different hydrogel samples using the same human donor. Data are shown as mean ± SD, and P-values from two-tailed unpaired Student's t-tests are indicated.

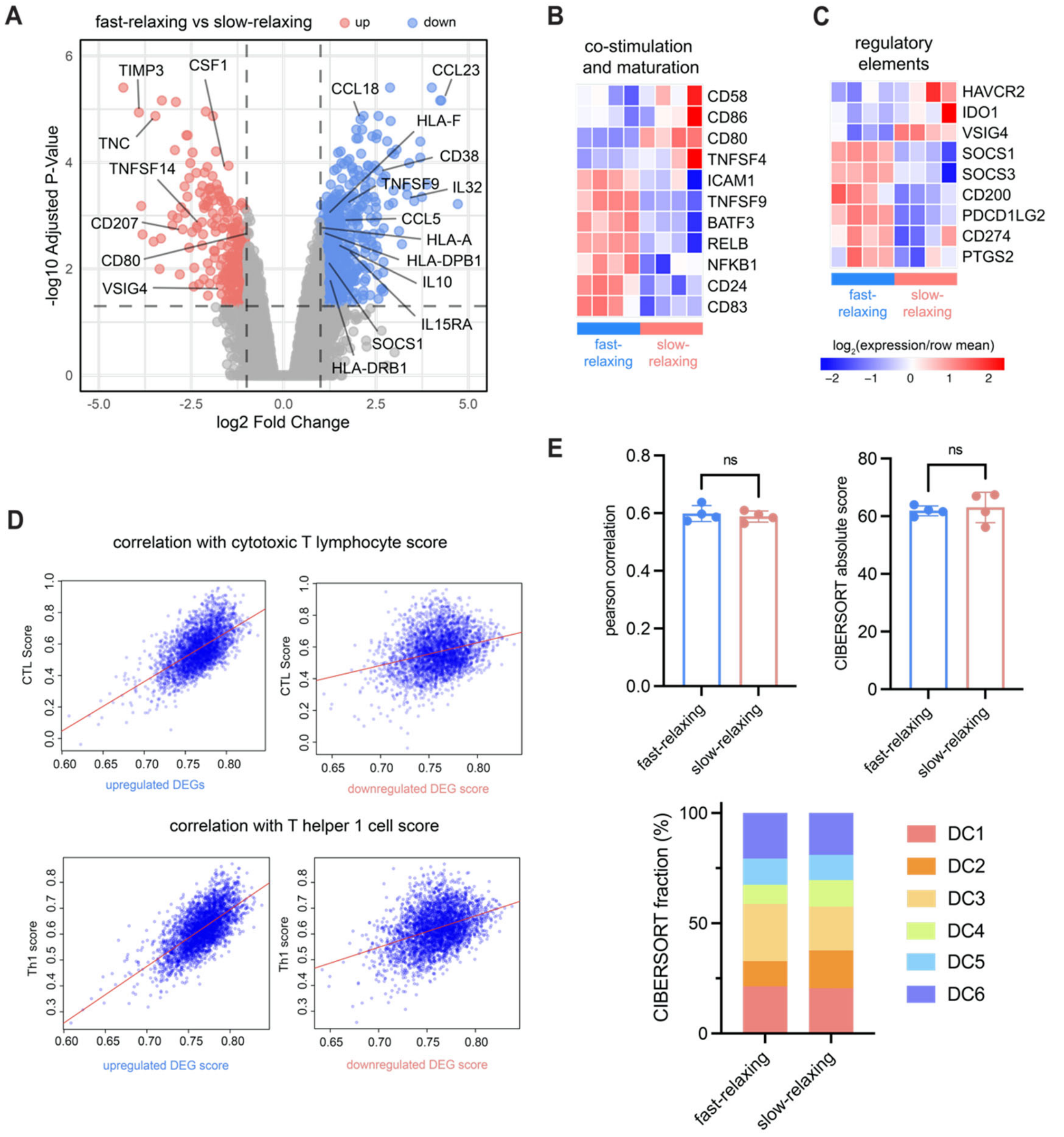


**Supplementary Figure S5 Related to Fig. 3. (A)** Volcano plot of differentially expressed genes between DCs from the fast-relaxing and slow-relaxing matrix. DEGs were filtered by log2 fold change > 1 and adjusted P-value < 0.05. Upregulated genes (enriched in fast-relaxing) are shown in blue and downregulated genes (enriched in slow-relaxing) are shown in red. **(B)** Heatmap of genes associated with co-stimulatory ligands and maturation signals. A differential enrichment of co-stimulatory molecules was observed between the two conditions, whereas the fast-relaxing matrix upregulated DC maturation signals such as BATF3, RELB, and NFKB1. **(C)** Heatmap of genes associated with regulatory elements at the immunological synapse. Consistent with the PD-L1 protein expression, the fast-relaxing matrix promoted gene expression of the regulatory elements such as PD-L1 (CD274) and PD-L2 (PDCD1LG2). On the contrary, T cell suppression signals Tim3 (HAVCR2), IDO1, and VSIG4 were enriched in the slow-relaxing matrix. **(D)** Scatter plots of ssGSEA scores calculated from the TCGA database. The y-axes are scores calculated from T cell signatures and the x-axes are scores from either the up- or down-regulated DEGs. Each patient sample from the database is shown as a blue dot (n = 3044) and the spearman correlation trendline is shown in red. **(E)** Pearson correlation and absolute score of CIBERSORT deconvolution. (top left) No statistical differences in the correlation between the bulk RNA sequencing from the mechanically educated DCs and the published single cell RNA sequencing from blood DCs were found. (top right) The absolute total CIBERSORT scores from all six subsets were not statistically different between the conditions, supporting the use of relative abundance for comparison. (bottom) The average CIBERSORT fractions from each DC subset from each condition. The majority of changes between distribution of subsets was observed in the DC2, DC3, and DC4 fractions. Data in A, B, C, and E are representative of n = 4 biological replicates from different hydrogels using the same human donor. Each column of the heatmap in B and C indicates a single biological replicate. Each dot in E indicates one replicate, and are shown as mean ± SD. P-values from two-tailed unpaired Student's t-tests are indicated.

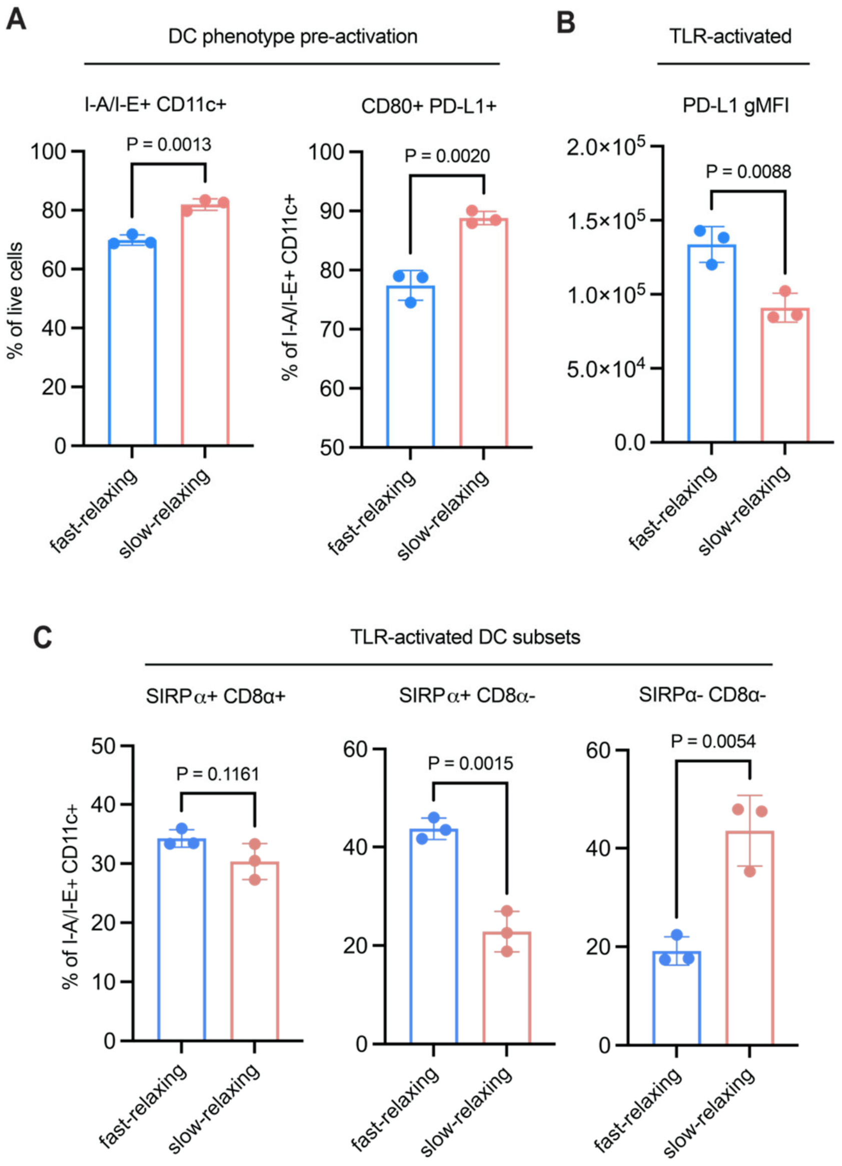


**Supplementary Figure S6 Related to Fig. 4. (A)** Mechanically educated murine DCs exhibited consistent trend as human DCs where the slow-relaxing matrix enriched the frequency of MHC-II+ CD11c+ CD80+ PD-L1+ DCs prior to TLR activation. **(B)** The fast-relaxing mDCs had higher PD-L1 expression after TLR activation, consistent with human DCs. **(C)** mDC subsets after TLR activation. The fast-relaxing DCs had higher frequency of SIRPα+ CD8α- cDC2 cells whereas the slow-relaxing DCs were enriched in a population lacking both conventional markers of cDC1 and cDC2, similar to the trend in the CD1c+ CD14+ CD163+ population and transcriptomic deconvolution of human DCs. Data are representative of n = 3 biological replicates from different hydrogel samples. Data are shown as mean ± SD, and P-values from two-tailed unpaired Student's t-tests are indicated.

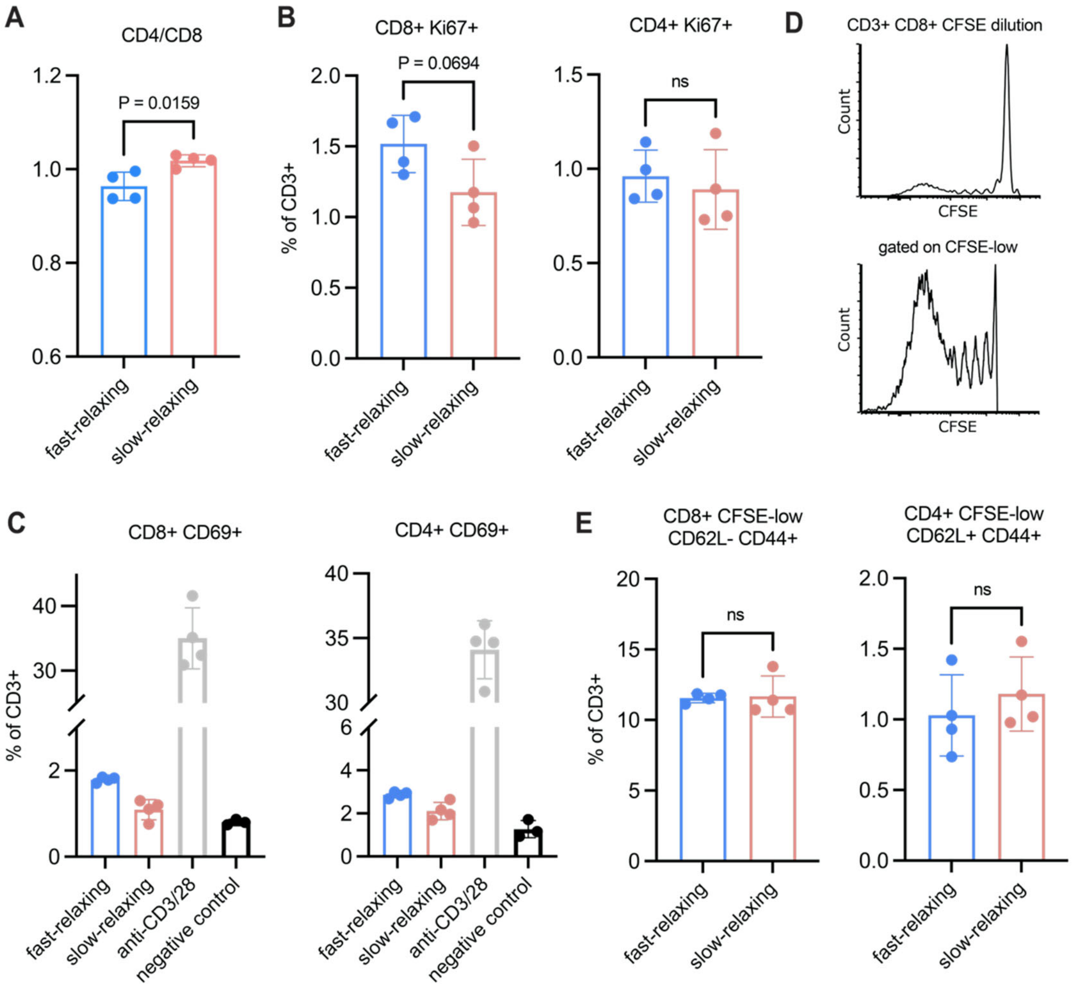


**Supplementary Figure S7 Related to Fig. 4. (A)** CD4/CD8 ratio of OVA-reactive T cells after 48 hours of co-culture with mechanically educated mDCs. The fast-relaxing DCs produced a CD8 biased response shown by the lower ratio and **(B)** higher frequency Ki67+ CD8+ T cells. **(C)** Positive control using plate-coated anti-CD3 and anti-CD28 showed high activation of CD69+ cells whereas the negative control showed the least amount of CD69+ cells in both CD4+ and CD8+ T cells. Data from the fast-relaxing and slow-relaxing conditions are the same as Fig. 4C. **(D)** Histogram of CFSE dilution of OVA-reactive T cells after 96 hours of co-culture with mechanically educated mDCs. **(E)** CFSE-low CD8+ T cells showed no statistical difference in effector memory T cell response and CFSE-low CD4+ T cells showed no statistical difference in central memory T cell response. Data are representative of n = 4 biological replicates. Data are shown as mean ± SD, and P-values from two-tailed unpaired Student's t-tests are indicated.

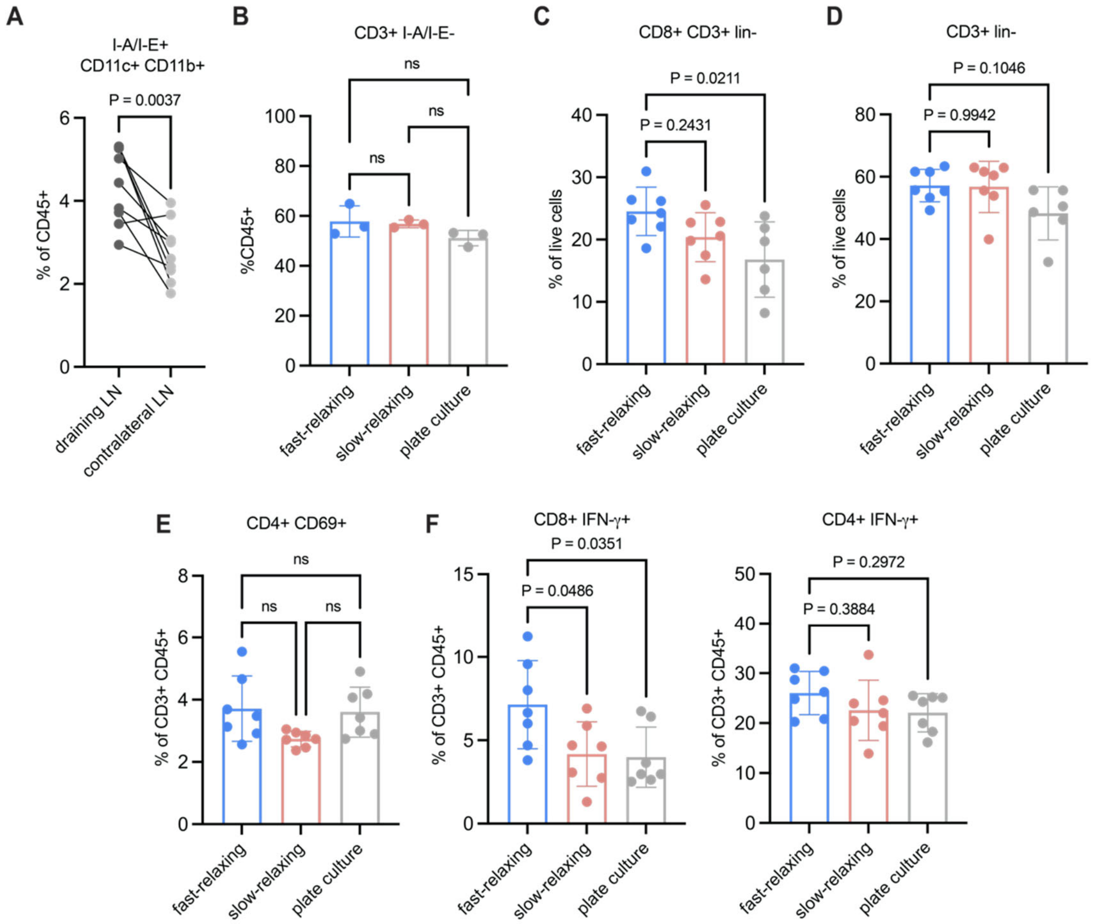


**Supplementary Figure S8 Related to Fig. 5. (A)** Paired biological replicate comparison of DC recruitment in draining lymph node and contralateral lymph node 24 hours after DC injection. The draining LN had higher frequency of MHC-II+ CD11c+ CD11b+ DCs, indicating an asymmetric immune response from the flank injection. **(B)** No statistical differences were observed in the abundance of dLN T cell population among the mechanically educated DC conditions 24 hours post injection. **(C)** 10 days after the DC injection, the fast-relaxing DCs induced a higher population of CD8+ T cells, but **(D)** no differences in the frequency of total T cells in the dLN. **(E)** No differences in the activation of CD69+ cells in the CD4+ compartment were observed upon peptide restimulation of the dLN cells. **(E, F)** Peptide restimulation produced a higher IFN-γ response in the CD8+ T cells in the dLNs primed by the fast-relaxing DCs.

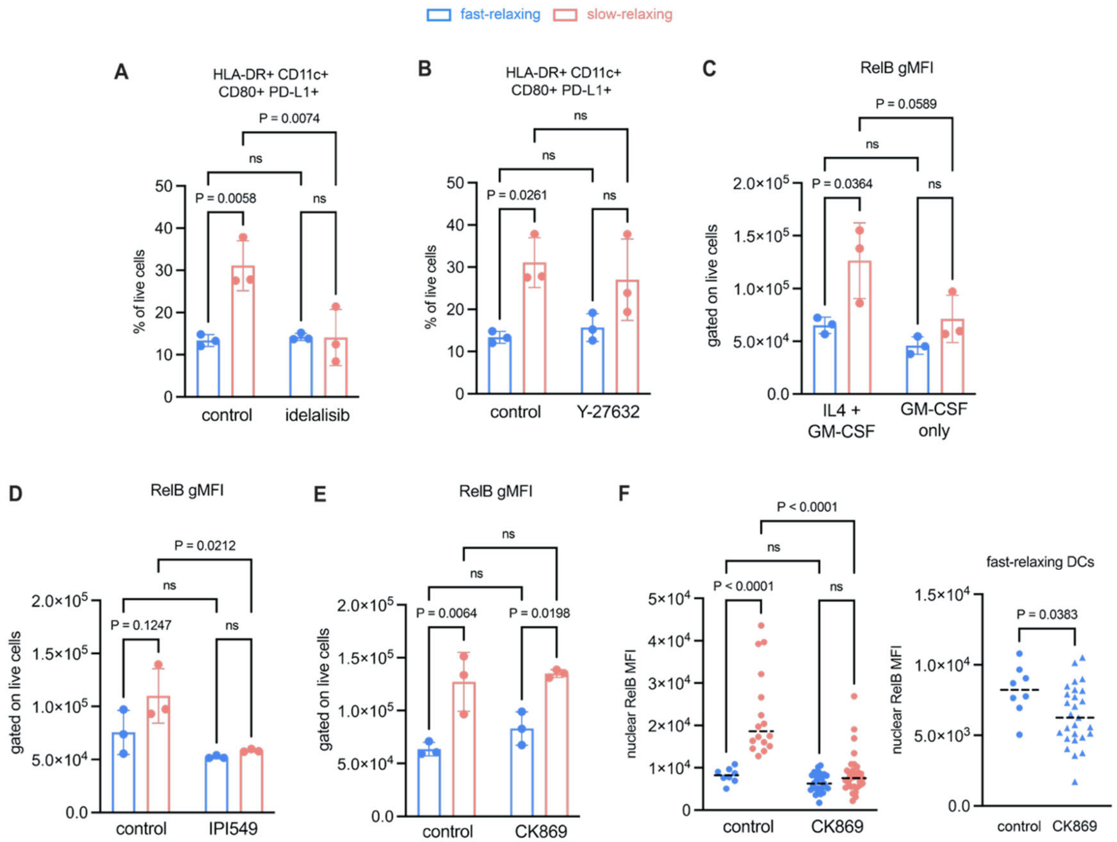


**Supplementary Figure S9 Related to Fig. 6. (A)** The slow-relaxing state was inhibited by a different PI3K inhibitor idelalisib targeting the delta isoform, consistent with IPI549 (gamma isoform). The control data are the same as Fig. 6C. **(B)** The slow-relaxing state was not perturbed by inhibiting Rho-ROCK pathway using Y-27632. The control data are the same as Fig. 6D. **(C)** High total RelB expression (quantified by intracellular flow cytometry) induced by the slow-relaxing matrix was dependent on the concerted cytokine signaling of IL-4 and GM-CSF. **(D)** Total RelB expression was decreased upon IPI540 inhibition in the slow-relaxing matrix. **(E)** Inhibition of Arp2/3 complex through CK869 had no impact on total RelB expression. However, **(F)** the nuclear expression upon Arp2/3 inhibition was decreased in both fast- and slow-relaxing matrix. Repeated control data are from the same experiment done with two different inhibitors.

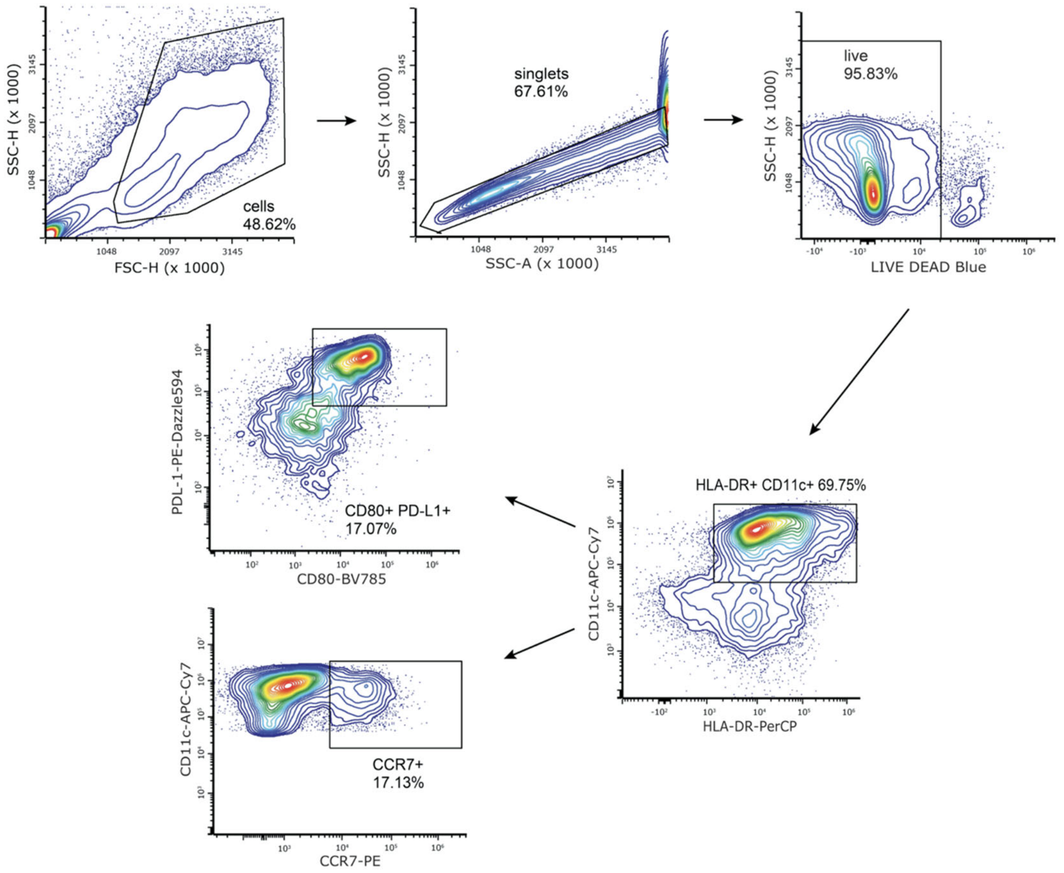


**Supplementary Figure S10.** Representative gating strategies of flow-cytometry. Live, single cells were gated prior to any analyses of marker expression. Gating was performed on unstained controls.

**Supplementary Table I. Hydrogel formulations**

| Hydrogel | Unmodified alginate (wt%) | Nb-modified alginate (wt%) | Tz-modified alginate (wt%) | Collagen (mg/ml) | Precipitated $CaCO_3$ (mM) |
|---|---|---|---|---|---|
| **Fast-relaxing** | 1 | 0 | 0 | 4 | 30 |
| **Slow-relaxing** | 0 | 0.8 | 0.2 | 4 | 10 |

**Supplementary Table II. Primary and secondary antibodies used in this study**

| Antibody | Specification | Dilution | Source |
|---|---|---|---|
| PerCP anti-human HLA-DR | Mouse monoclonal | 1:100 | Biolegend (307628) |
| Brilliant Violet 510 anti-human CD1c | Mouse monoclonal | 1:100 | Biolegend (331534) |
| APC/Cyanine7 anti-human CD11c | Mouse monoclonal | 1:100 | Biolegend (337218) |
| Alexa Fluor 647 anti-human CD14 | Mouse monoclonal | 1:100 | Biolegend (301812) |
| Brilliant Violet 605 anti-human CD163 | Mouse monoclonal | 1:100 | Biolegend (333615) |
| Brilliant Violet 785 anti-human CD80 | Mouse monoclonal | 1:100 | Biolegend (305238) |
| PE-Dazzle594 anti-human PD-L1 (CD274) | Mouse monoclonal | 1:100 | Biolegend (329371) |
| PE anti-human CCR7 (CD197) | Mouse monoclonal | 1:100 | Biolegend (353203) |
| PE/Cyanine5 anti-human CD86 | Mouse monoclonal | 1:100 | Biolegend (305407) |
| Alexa Fluor 594 anti-human CD3 | Mouse monoclonal | 1:100 | Biolegend (300446) |
| Pacific blue anti-human CD40 | Mouse monoclonal | 1:100 | Biolegend (334320) |
| Pacific blue anti-mouse I-A/I-E | Rat monoclonal | 1:100 | Biolegend (107620) |
| Alexa Fluor 700 anti-mouse CD11c | Rat monoclonal | 1:100 | Biolegend (117320) |
| Alex Fluor 488 anti-mouse CD172a (SIRPalpha) | Rat monoclonal | 1:100 | Biolegned (144024) |
| APC anti-mouse PD-L1 (CD274) | Rat monoclonal | 1:100 | Biolegend (124312) |
| PE/Cyanine7 anti-mouse CD80 | Rat monoclonal | 1:100 | Biolegend (104734) |
| PerCP/Cyanine5.5 anti-mouse CD8a | Rat monoclonal | 1:100 | Biolegend (155013) |
| PE anti-mouse H-2Kb bound to SIINFEKL | Mouse monoclonal | 1:100 | Biolegend (141603) |
| PerCP anti-mouse CD3 | Rat monoclonal | 1:100 | Biolegend (100287) |
| Alexa Fluor 594 anti-mouse/human Ki-67 | Rat monoclonal | 1:100 | Biolegend (151213) |
| PE anti-mouse CD62L | Rat monoclonal | 1:100 | Biolegend (161203) |
| APC/Cyanine7 anti-mouse CD69 | Rat monoclonal | 1:100 | Biolegend (104525) |
| APC anti-mouse CD4 | Rat monoclonal | 1:100 | Biolegend (100411) |
| FITC anti-mouse CD45 | Rat monoclonal | 1:100 | Biolegend (103108) |
| Alexa Fluor 700 anti-mouse IFN-gamma | Rat monoclonal | 1:100 | Biolegend (505823) |

| | | | |
|---|---|---|---|
| PE/Cyanine5 anti-mouse/human CD44 | Rat monoclonal | 1:100 | Biolegend (103009) |
| Pacific Blue anti-mouse CD19 | Rat monoclonal | 1:100 | Biolegend (152415) |
| RelB (D7D7W) | Rabbit monoclonal | 1:800 | Cell Signaling Technologies (10544) |
| Alex Fluor 488 anti-rabbit IgG | Goat Polyclonal | 1:500 | Invitrogen (A-11034) |

**Supplementary Table III. P-value table**

| Figure | Statistical Test | Comparison | P-value |
|---|---|---|---|
| Fig. 1D | Two-tailed Student t-test | FR vs SR | 0.5985 |
| Fig. 1F HLA-DR | Two-tailed Student t-test | FR vs SR | 0.0031 |
| Fig. 1F CD80 | Two-tailed Student t-test | FR vs SR | 0.0158 |
| Fig. 1F CD1c | Two-tailed Student t-test | FR vs SR | 0.0127 |
| Fig. 2B | Two-tailed Student t-test | FR vs SR | 0.0056 |
| Fig. 2C CD86 | Two-tailed Student t-test | FR vs SR | 0.0276 |
| Fig. 2C CD80 | Two-tailed Student t-test | FR vs SR | 0.0057 |
| Fig. 2C PD-L1 | Two-tailed Student t-test | FR vs SR | 0.0039 |
| Fig. 2D IL-6 | Two-tailed Student t-test | FR vs SR | 0.0005 |
| Fig. 2D TNF-a | Two-tailed Student t-test | FR vs SR | < 0.0001 |
| Fig. 2D IL-10 | Two-tailed Student t-test | FR vs SR | 0.0008 |
| Fig. 2E CD4 | Two-tailed Student t-test | FR vs SR | 0.0081 |
| Fig. 2E CD8 | Two-tailed Student t-test | FR vs SR | 0.0027 |
| Fig. 3G DC1 | Two-tailed Student t-test | FR vs SR | 0.5527 |
| Fig. 3G DC2 | Two-tailed Student t-test | FR vs SR | 0.0019 |
| Fig. 3G DC3 | Two-tailed Student t-test | FR vs SR | 0.0025 |
| Fig. 3G DC4 | Two-tailed Student t-test | FR vs SR | 0.0196 |
| Fig. 3H | Two-tailed Student t-test | FR vs SR | 0.0002 |
| Fig. 4B H2Kb | Two-tailed Student t-test | FR vs SR | 0.0045 |
| Fig. 4B CD80 | Two-tailed Student t-test | FR vs SR | 0.1498 |
| Fig. 4C CD8 | Two-tailed Student t-test | FR vs SR | 0.0013 |
| Fig. 4C CD4 | Two-tailed Student t-test | FR vs SR | 0.0128 |
| Fig. 4D CD8 | Two-tailed Student t-test | FR vs SR | 0.0426 |
| Fig. 4D CD8 CFSE | Two-tailed Student t-test | FR vs SR | 0.0284 |
| Fig. 4D CD4 | Two-tailed Student t-test | FR vs SR | 0.0017 |
| Fig. 4D CD4 CFSE | Two-tailed Student t-test | FR vs SR | 0.0003 |
| Fig. 5B | One-way ANOVA & Tukey | FR vs SR | 0.9846 |
| Fig. 5B | One-way ANOVA & Tukey | FR vs Plate | 0.4555 |
| Fig. 5B | One-way ANOVA & Tukey | SR vs Plate | 0.3778 |
| Fig. 5C tetramer | One-way ANOVA & Tukey | FR vs SR | 0.0477 |
| Fig. 5C tetramer | One-way ANOVA & Tukey | FR vs Plate | 0.2475 |
| Fig. 5C tetramer | One-way ANOVA & Tukey | SR vs Plate | 0.6936 |
| Fig. 5C CD62L- | One-way ANOVA & Tukey | FR vs SR | 0.0032 |
| Fig. 5C CD62L- | One-way ANOVA & Tukey | FR vs Plate | 0.0039 |
| Fig. 5C CD62L- | One-way ANOVA & Tukey | SR vs Plate | 0.9973 |
| Fig. 5D CD4 Ki67 | One-way ANOVA & Tukey | FR vs SR | 0.0177 |

| | | | |
|---|---|---|---|
| Fig. 5D CD4 Ki67 | One-way ANOVA & Tukey | FR vs Plate | 0.0095 |
| Fig. 5D CD4 Ki67 | One-way ANOVA & Tukey | SR vs Plate | 0.9545 |
| Fig. 5D CD8 Ki67 | One-way ANOVA & Tukey | FR vs SR | 0.0138 |
| Fig. 5D CD8 Ki67 | One-way ANOVA & Tukey | FR vs Plate | 0.0064 |
| Fig. 5D CD8 Ki67 | One-way ANOVA & Tukey | SR vs Plate | 0.9311 |
| Fig. 5D CD8 CD69 | One-way ANOVA & Tukey | FR vs SR | 0.0217 |
| Fig. 5D CD8 CD69 | One-way ANOVA & Tukey | FR vs Plate | 0.4901 |
| Fig. 5D CD8 CD69 | One-way ANOVA & Tukey | SR vs Plate | 0.1979 |
| Fig. 5E | One-way ANOVA & Tukey | FR vs SR | 0.0482 |
| Fig. 5E | One-way ANOVA & Tukey | FR vs Plate | 0.8021 |
| Fig. 5E | One-way ANOVA & Tukey | SR vs Plate | 0.1581 |
| Fig. 6A volume | Two-tailed Student t-test | FR vs SR | 0.0102 |
| Fig. 6A sphericity | Two-tailed Student t-test | FR vs SR | 0.0371 |
| Fig. 6A F-actin | Two-tailed Student t-test | FR vs SR | 0.0171 |
| Fig. 6B | Two-way ANOVA & Tukey | FR IL4 vs SR IL4 | 0.0035 |
| Fig. 6B | Two-way ANOVA & Tukey | FR IL4 vs FR GM | 0.1122 |
| Fig. 6B | Two-way ANOVA & Tukey | FR GM vs SR GM | 0.5584 |
| Fig. 6B | Two-way ANOVA & Tukey | SR IL4 vs SR GM | 0.0008 |
| Fig. 6C | Two-way ANOVA & Tukey | FR con vs SR con | 0.0006 |
| Fig. 6C | Two-way ANOVA & Tukey | FR con vs FR IPI | 0.9972 |
| Fig. 6C | Two-way ANOVA & Tukey | FR IPI vs SR IPI | 0.8916 |
| Fig. 6C | Two-way ANOVA & Tukey | SR con vs SR IPI | 0.0016 |
| Fig. 6D | Two-way ANOVA & Tukey | FR con vs SR con | < 0.0001 |
| Fig. 6D | Two-way ANOVA & Tukey | FR con vs FR CK | < 0.0001 |
| Fig. 6D | Two-way ANOVA & Tukey | FR CK vs SR CK | 0.0072 |
| Fig. 6D | Two-way ANOVA & Tukey | SR con vs SR CK | 0.2256 |
| Fig. 6E | Two-tailed Student t-test | FR vs SR | < 0.0001 |
